# LINC complex alterations are a hallmark of sporadic and familial ALS/FTD

**DOI:** 10.1101/2024.03.08.584123

**Authors:** Riccardo Sirtori, Michelle Gregoire, Emily Potts, Alicia Collins, Liviana Donatelli, Claudia Fallini

## Abstract

Amyotrophic lateral sclerosis (ALS) is a neurodegenerative disorder that primarily affects motor neurons, leading to progressive muscle weakness and loss of voluntary muscle control. While the exact cause of ALS is not fully understood, emerging research suggests that dysfunction of the nuclear envelope (NE) may contribute to disease pathogenesis and progression. The NE plays a role in ALS through several mechanisms, including nuclear pore defects, nucleocytoplasmic transport impairment, accumulation of mislocalized proteins, and nuclear morphology abnormalities. The LINC complex is the second biggest multi-protein complex in the NE and consists of the SUN1/2 proteins spanning the inner nuclear membrane and Nesprin proteins embedded in the outer membrane. The LINC complex, by interacting with both the nuclear lamina and the cytoskeleton, transmits mechanical forces to the nucleus regulating its morphology and functional homeostasis. In this study we show extensive alterations to the LINC complex in motor and cortical iPSC-derived neurons and spinal cord organoids carrying the ALS causative mutation in the *C9ORF72* gene (C9). Importantly, we show that such alterations are present *in vivo* in a cohort of sporadic ALS and C9-ALS postmortem spinal cord and motor cortex biopsies. We also found that LINC complex disruption strongly correlated with nuclear morphological alterations occurring in ALS neurons, independently of TDP43 mislocalization. Altogether, our data establish morphological and functional alterations to the LINC complex as important events in ALS pathogenic cascade, making this pathway a possible target for both biomarker and therapy development.

## Introduction

Amyotrophic lateral sclerosis (ALS) is the most common adult-onset motor neuron disease, characterized by progressive muscle atrophy and weakness that eventually results in respiratory failure and death. Over 90% of ALS cases are not linked to clear familial inheritance and have no defined cause (i.e., sporadic ALS or sALS). The remaining 10% of ALS cases are associated with pathogenic mutations in one of more than 30 different genes^1^. One of the most common genetic causes for both ALS and frontotemporal dementia (FTD), a closely related neurodegenerative condition that shares both genetic and pathological signs with ALS, is a *GGGGCC* hexanucleotide repeat expansion (HRE) in the *C9ORF72* gene (C9-ALS)^2^. This mutation accounts for about 40% of familial and 7% of sporadic ALS cases^3^. Although the molecular determinants of neuronal degeneration in ALS/FTD remain generally unclear, protein aggregation, most prominently of the RNA-binding protein TDP-43^4^, and alterations to nuclear homeostasis^5^ have been accepted as common pathomechanisms underlying most familial and sporadic forms of the disease. In fact, alterations to the nuclear lamina and to the nuclear pore complex (NPC), the largest multiprotein assembly encompassing the inner and outer nuclear membranes, have been widely reported in several *in vivo* and *in vitro* models of ALS/FTD, including postmortem patients’ tissue. Of note, we have recently shown that mechanical strain on the nucleus, exerted by the cytoskeleton via the linker of nucleoskeleton and cytoskeleton (LINC) complex, can lead to NPC injury, ruptures of the nuclear envelope (NE), and accumulation of DNA damage^6^, significantly contributing to disease pathology. One of the cell’s main mechanosensor, the LINC complex is the second largest transmembrane complex present in the NE. While its modular protein composition is variable in different cell types and conditions, the LINC complex generally consists of a trimer of SUN proteins (SUN1 and SUN2) spanning the inner nuclear membrane, connected to a trimer of Nesprin proteins (Nesprin 1-4) embedded in the outer nuclear membrane^7^. The cytoplasmic domains of Nesprins bind to different cytoskeletal components, while their intermembrane Klarsicht, Anc-1, Syne homology (KASH) domains bind to SUN proteins, which are in turn bound to the nuclear lamina and NPCs^8^. Because of these multiple interactions, the LINC complex effectively couples the cytoskeleton to NPCs and the nuclear lamina, resulting in nuclear exposure to cytoskeletal forces^9^. These connections act as a fundamental determinant of nuclear morphology and size, chromatin organization, and nucleolar arrangement ^10,11^.

Mutations in components of the LINC complex are associated with several human diseases, including laminopathies^12^, skeletal muscle diseases^13^, neurodevelopmental diseases^14,15^, neurodegenerative conditions such as cerebellar ataxia with motoneuronal involvement^16^ and juvenile ALS^17,18^. In addition, reduced nuclear localization of SUN1 protein was recently described in sALS iPSCs-derived neurons *in vitro*^19^, suggesting that this complex may play a relevant role in ALS/FTD pathogenesis. In this study, we show that multiple protein components of the LINC complex are misregulated in different *in vitro* ALS models, as well as in patient’s postmortem tissue. Furthermore, we found that relative nuclear and nucleolar sizes are significantly reduced in sALS and C9-ALS neurons. Interestingly, nuclear and nucleolar shrinkage occurs even in the absence of common ALS pathological markers but correlates strongly with the degree of disruption of the LINC complex, suggesting that this pathological phenotype depends on the LINC complex and may precede TDP-43 mislocalization as an early sign of cell dysfunction.

## RESULTS

### 1. The LINC complex is altered in C9-ALS iPSC-derived motor and cortical neurons

To determine whether alterations to the LINC complex play a relevant role in ALS/FTD pathogenesis, we investigated the overall levels and cellular distribution of the main LINC protein components, namely SUN1, SUN2, Nesprin1 (NESP1), and Nesprin2 (NESP2). For this study, we first focused on iPSC-derived motor neurons (iMNs) carrying mutations in the *C9ORF72* locus, as this mutation has already been extensively implicated in nuclear envelope disfunctions ^20,21,22,23^. iMNs were derived from two pairs of isogenic iPSCs where the patient’s derived C9 HRE had been corrected using CRISPR/Cas9 technology (**Supplementary Table 1**). Immunostaining with neuronal and MN-specific markers including NeuN, MAP2 and ISL1 confirmed that both the patient mutant lines and their isogenic control lines generated ISL1-positive iMNs with a differentiation with similar efficiency (**Supplementary Figure 1**). Six weeks after differentiation induction, we found that the nuclear levels of all SUN and Nesprin proteins tested were severely reduced in C9 MNs compared to control isogenic neurons (**Figure 1** and **Supplementary Figure 2**). WB analyses also showed an overall reduction in the whole-cell levels of SUN2 (**Figure 1c**) but not SUN1 (**Supplementary Figure 2f**). However, we observed a reduction of SUN1 nucleo-cytoplasm ratio in C9 compared to control isogenic MNs, possibly accounting for this discrepancy (**Supplementary Figure 2c**).

**Figure 1.**
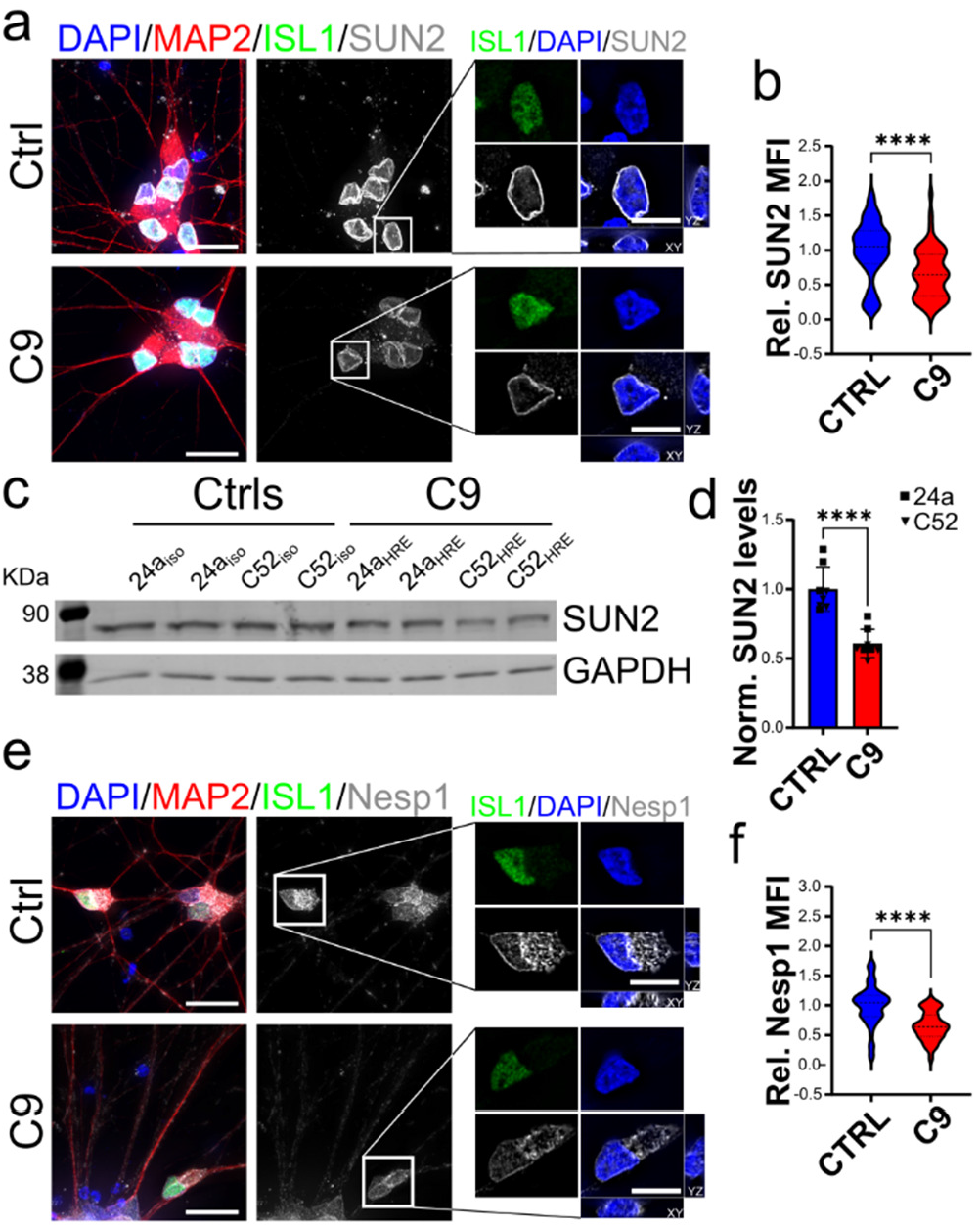
LINC complex disruption in C9 iPSCs-derived iMNs. **a-b**. Representative images of SUN2 (***a***, *grays*) and Nesprin1 (***e***, Nesp1, *grays*) expression in C9 and isogenic control (CTRL) iMNs. Nuclei were identified by DAPI staining (*blue*), while MAP2 (*red*) and ISL1 (*green*) were used as neuronal and motoneuronal markers. The white boxes indicate the neurons enlarged in the panels on the right. Scale bars: 20μm in main panels, 10μm in zoomed-in images. The relative quantification of the nuclear mean fluorescence intensity (MFI) for both SUN2 (***b***) and Nesprin1 (***f***) shows a significant reduction in the abundance of each protein in C9 iMNs compared to isogenic controls (Mann-Whitney *t* test, n= 57 and 52 for CTRL and C9-ALS neurons respectively from 4 independent differentiations for SUN2, n= 64 and 40 for CTRL and C9 neurons respectively from 4 independent differentiations for Nesprin1, **** *p* < 0.0001). **c-d**. Representative western blot (WB) and quantification of SUN2 levels relative to GAPDH expression shows a significative reduction of total SUN2 levels in both C9 isogenic lines (24a and C52) compared to isogenic counterparts (n=8 independent experiments for both C9 and controls; Student’s *t* test, **** *p* < 0.0001). For all, bars are mean and SEM, while violin plots show the distribution of the data with dashed lines indicating median and quartiles.

Since the hexanucleotide expansion in *C9ORF72* is also causative for frontotemporal lobar degeneration (FTLD), accounting for about 25% of familial and 5% of sporadic cases^24^, we decided to expand our initial observations in C9-ALS iMNs to cortical neurons differentiated from the same iPSC lines as above using the previously published i^3^ method ^25^. Two weeks after the induction of differentiation, all the cells were positive for the neuronal markers NeuN and MAP2, displayed glutamate receptor surface expression, and spontaneous electrical activity (**Supplementary Figure 3**). Under these conditions, we found that the nuclear levels of all the LINC proteins tested were severely reduced in C9 iPSC-derived cortical neurons (i^3^CNs), similarly to what observed in the iMNs (**Figure 2** and **Supplementary Figure 4**). This decrease, quantified by IF assays, was associated to a significant reduction in their whole cell levels, quantified by WB for SUN1 and SUN2 (**Figure 2f** and **Supplementary Figure 4f-g**). We further confirmed these observations by qualitatively scoring nuclei based on SUN or NESP proteins distribution at the NE. While the linear profile of SUN and NESP proteins fully matched the DAPI or Lamin B (LMNB) profiles in isogenic i^3^CNs, mutant neurons were characterized by a higher frequency of smaller intensity peaks irregularly distributed across the whole cell (**Figures 2b-d, 2i-k**, and **Supplementary Figures 4b-d, 4i-k**). Altogether, these observations suggest that disruption of the LINC complex may be an early phenomenon occurring in C9-ALS.

**Figure 2.**
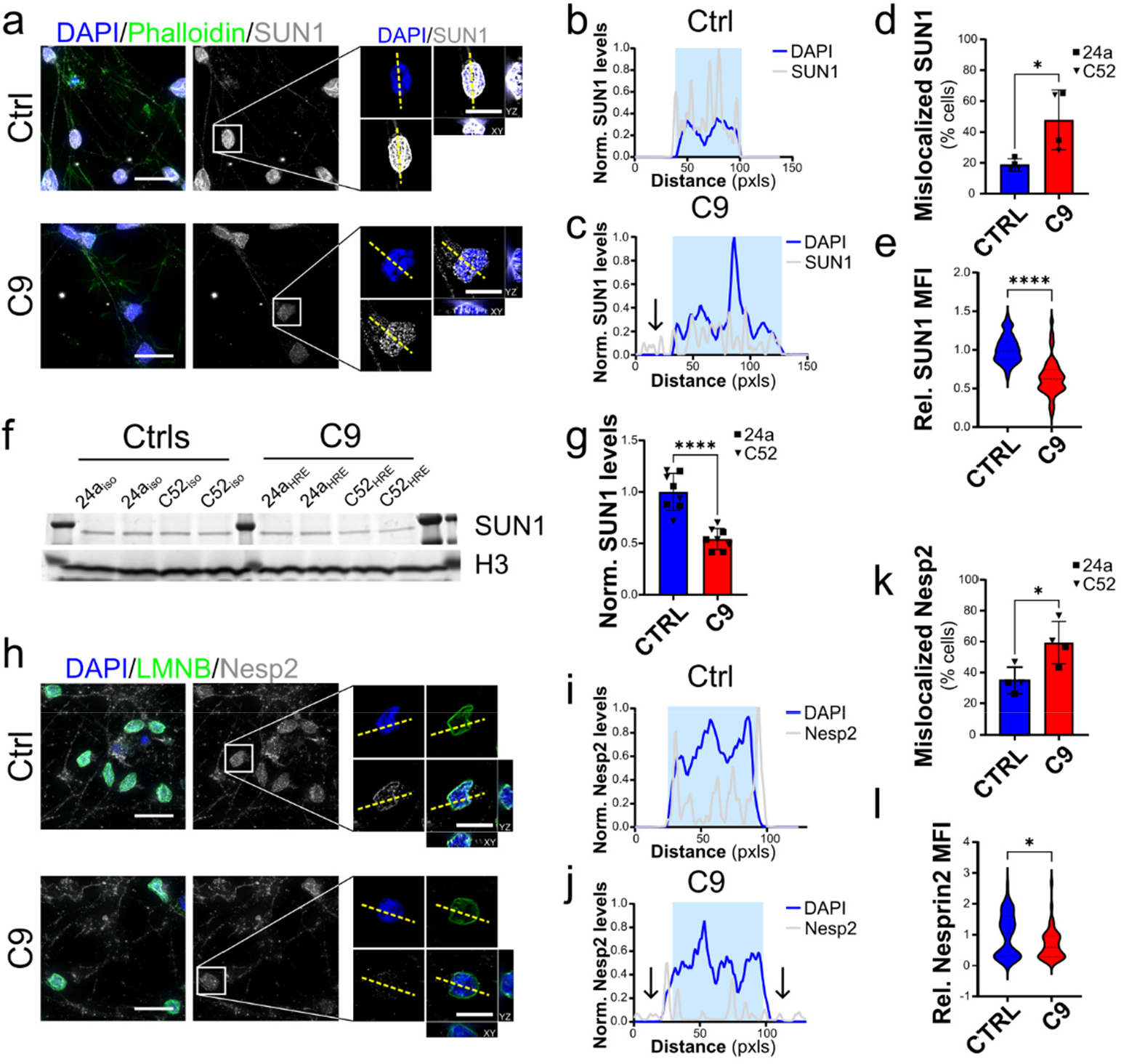
LINC complex disruption in C9-ALS i^3^CNs. **a**. Representative images of SUN1 (*grays*) in C9-ALS and control i^3^CNs. DAPI (*blue*) identified the nuclei, while Phalloidin (*green*) labeled the actin cytoskeleton. The white boxes indicate the neurons enlarged in the panels on the right. **b-d**. Plots of line profiles of SUN1 and DAPI intensities normalized to the max intensity in control cells. Mutant cells display frequent mislocalization of SUN1 to either the nucleoplasm or cytoplasm (arrows), quantified in *d*. The yellow dashed lines in *a* indicate the lines used for the profile plots (Mann-Whitney *t* test, n= 4, \**p*<0.05). **e**. The quantification of the nuclear mean fluorescence intensity (MFI) of SUN1 in C9 i^3^CNs shows a significant reduction in its abundance relative to isogenic controls (Mann-Whitney *t* test, n= 46 CTRL and C9 neurons respectively from 4 independent differentiations, \*\*\*\**p*<0.0001). **f-g**. Representative western blot (WB) and quantification of SUN1 levels relative to Histone 3 (H3) expression shows a significative reduction of total SUN1 levels in C9 lines compared to isogenic counterparts (n = 8 independent experiments for both 24a and C52 iPSC isogenic pairs; Student’s *t* test, **** *p* < 0.0001). **h**. Representative images of Nesprin2 (Nesp2) staining pattern in i^3^CNs from C9 and Ctrl iPSC lines. DAPI (*blue*) identified the nuclei, while LaminB (LMNB, *green*) labeled the nuclear lamina. The white boxes indicate the neurons enlarged in the panels on the right. **i-k**. Plots of line profiles of Nesprin2 and DAPI intensities normalized to the max intensity in control cells. Mutant cells display frequent mislocalization of Nesprin2 to either the nucleoplasm or cytoplasm (arrows), quantified in *k*. The yellow dashed lines in *h* indicate the lines used for the profile plots (Student’s *t* test, n= 4, \**p*<0.05). **l**. The quantification of the Nesprin2 relative nuclear MFI shows a significant reduction in its abundance in C9 i^3^CNs compared to isogenic controls (Mann-Whitney *t* test, n= 31 and 38 for CTRL and C9 neurons from 4 independent differentiations, * *p* < 0.05). Scale bars: 20μm in the main panels, 10μm in zoomed-in images. For all, bars are mean and SEM, while violin plots show the distribution of the data with dashed lines indicating median and quartiles.

### 2. SUN1 and SUN2 levels are reduced in C9-ALS iPSC-derived spinal cord organoids

Given the importance of the LINC complex in sensing mechanical properties of the cell’s intracellular and extracellular environment, which are not fully reproduced in a 2D cell culture system, we decided to assess LINC complex alterations in spinal cord organoids, which were generated using established protocols ^26^. As expected, after 25 days of differentiation we could clearly identify NeuN-positive neurons and ISL1-positive MNs within each organoid, while GFAP-positive cells appeared around day 40 (**Supplementary Figure 5**). When we quantified SUN1 and SUN2 protein distribution and NE abundance by IF, we found a significant reduction of their nuclear levels in ISL1-positive C9 iMNs at day 50 (**Figure 3g** and **3n**), which was also coupled by the alteration of their distribution, as shown by the analysis of their line profiles. In fact, we found that mutant neurons were characterized by a loss of high-intensity peaks at the periphery of the DAPI profile and an increase in intranuclear peaks (**Figure 3b-e**, and **3i-l**). This was confirmed by the blind assessment of SUN1 and SUN2 frequency of mislocalization from the NE (**Figure 3f, m**). An overall reduction of both SUN1 and SUN2 protein levels was also observed by WB using whole protein extracts of 120-day-old organoids (**Figure o-r**). Altogether these results confirm the impairment of the LINC complex in a 3D environment such as C9 iPSCs-derived spinal cord organoids.

**Figure 3.**
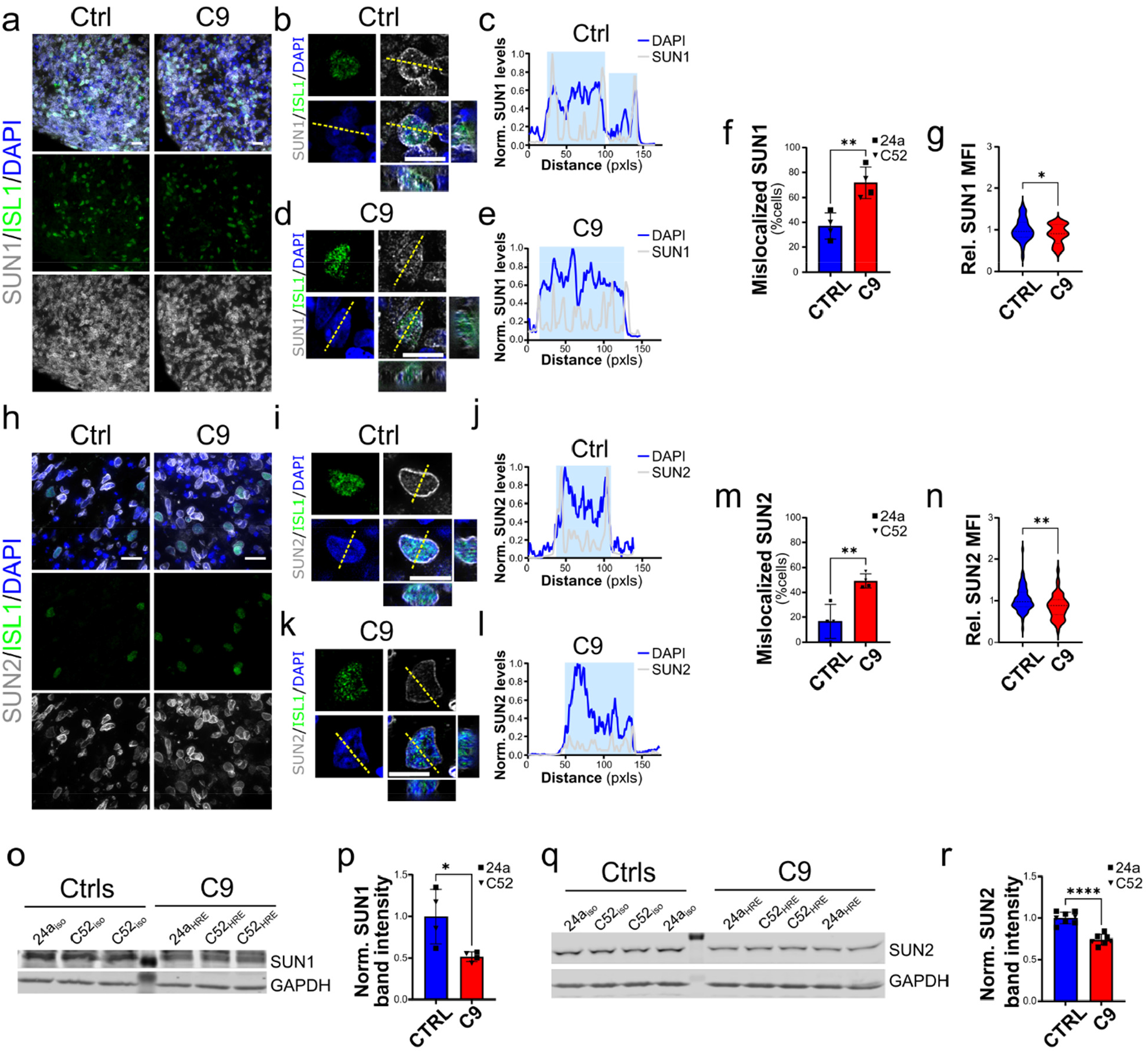
SUN1 and SUN2 alterations in spinal organoid iMNs. Representative images of SUN1 (***a***, *grays*) and SUN2 (***h***, *grays*) staining in spinal organoids. Islet1 (ISL1, *gree*n) expression was used to identify iMNs, while DAPI (*blue*) labeled the nuclei. **b-f**. Representative images and line profiles of SUN1 distribution in iMNs from control (*b-c*) and C9 mutant (*d-e*) organoids show a disruption in its localization at the NE, which was more frequently observed in iMNs from mutant organoids (quantified in *f*; Student’s *t* test, n=4, \*\**p*<0.01). The yellow dashed lines in *a* indicate the lines used for the profile plots. **g**. Quantitative analysis of SUN1 nuclear levels shows a significant reduction in C9 organoids compared to isogenic control (Student’s *t* test, n=71 and 31 from 4 independent experiments, \**p*<0.05). **i-m**. Representative images and line profiles of SUN2 distribution in iMNs from control (*i-j*) and C9 mutant (*k-l*) organoids show a disruption in its localization at the NE, which was more frequently observed in iMNs from mutant organoids (quantified in m; Student’s *t* test, n= 4, \*\**p*<0.01). **n**. Quantitative analysis of SUN2 nuclear levels shows a significant reduction in C9 organoids compared to isogenic control (Student’s *t* test, n=71 and 47 from 4 independent experiments, \*\**p*<0.01). **o-r**. Representative blots and quantification of SUN1 (o-p) and SUN2 (q-r) levels from whole lysates of spinal organoids from C9 and control iPSCs shows a significant reduction in their overall levels (Student’s *t* test, n=4 in *p*, n=7 and 8 in *r*, \**p*<0.05, \*\*\*\**p*<0.0001). GAPDH was used as loading control. For all, bars are mean and SEM, while violin plots show the distribution of the data with dashed lines indicating median and quartiles. Scale bars: 20μm in a and h, 10μm in b, d, i, and k.

### 3. LINC complex is disrupted in motor neurons of sporadic and C9-ALS spinal cord

Since our data in 2D and 3D culture systems of both cortical and motor neurons strongly suggested that disruption to the LINC complex may be relevant to ALS/FTD disease pathogenesis, we decided to confirm such findings using patient-derived *postmortem* tissues. We first investigated the distribution of SUN and NESP proteins in the spinal cord of 5 sALS and 3 familial C9-ALS biopsies compared to 5 healthy controls (**Supplementary Table 2** and **3**). These patients were all diagnosed with definite ALS and presented with a combination of upper and lower motor neuron signs and concomitant pathology. Only one C9-ALS patient also presented signs and symptoms related to FTD pathology. Given that SUN and NESP immunoreactivity creates a distinct ring around the nuclei of neurons, we scored morphologically defined motor neurons in the ventral horn blindly, based on the presence of a complete perinuclear ring at their equatorial plan (**Figure 4** and **Supplementary Figure 6**). We found a significant higher percentage of MNs presenting an altered or undetectable nuclear localization of both SUN and NESP proteins in MNs of sALS and C9-ALS compared with healthy controls, while no significant difference was noted between sALS and C9-ALS (**Figure 4g-h, Supplementary Figure 6g-h**).

**Figure 4.**
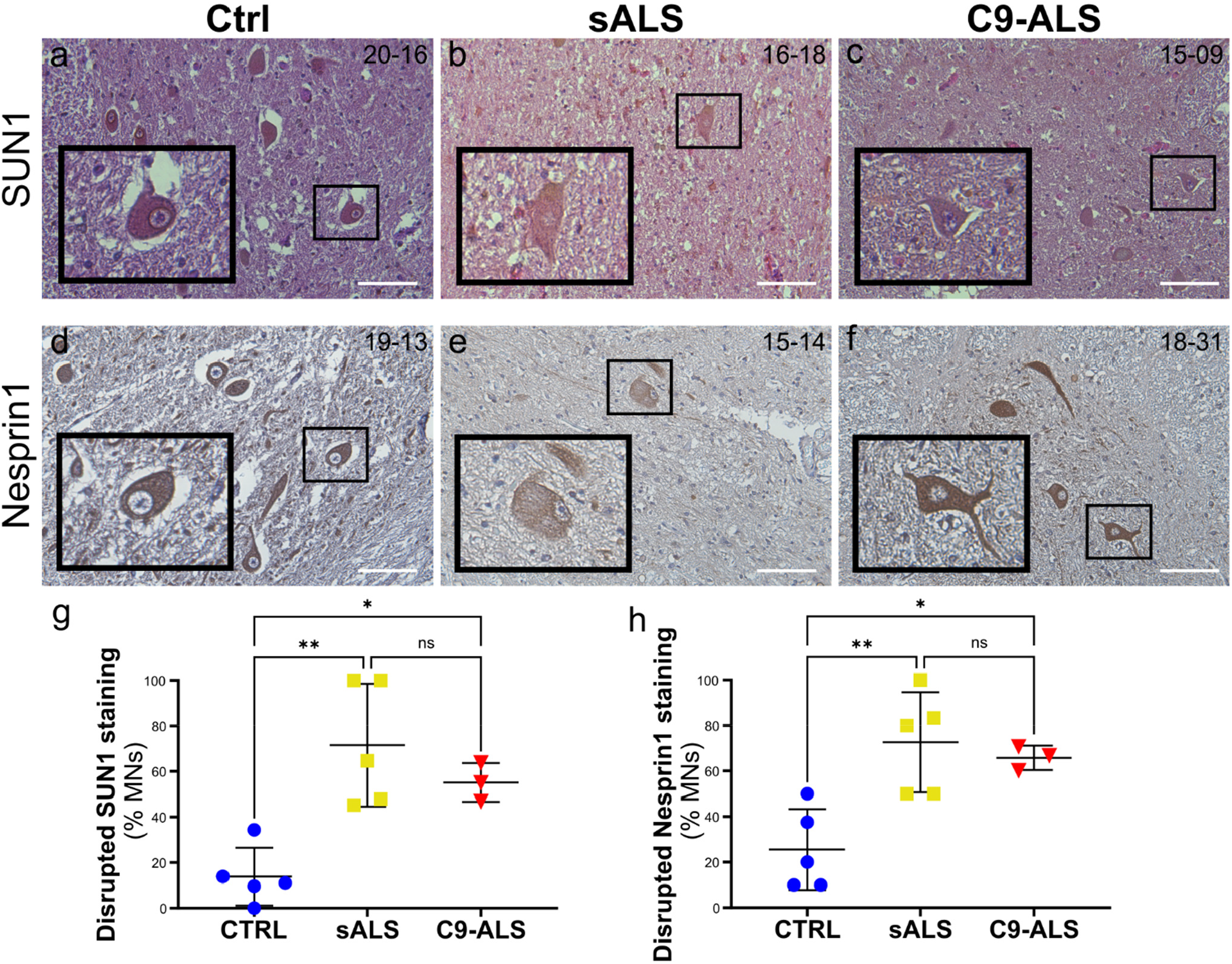
LINC complex disruption in sALS and C9-ALS spinal cord *postmortem* biopsies. **a-f**. Spinal cord sections from control (*a, d*), sALS (*b, e*) and C9-ALS (*c, f*) patients were stained with antibodies specific for SUN1 (*a-c*) and Nesprin1 (*d-f*). Hematoxylin and eosin counterstains were used to identify the nucleus and cytoplasm, respectively. The black boxes identify the motor neurons enlarged in the insets. Scale bars: 100μm. **g-h**. The frequency of disrupted NE staining for both SUN1 (*g*) and Nesprin1 (*h*) was quantified blindly in at least two sections from each patient’s tissue. A significant increase in the percentage of cells with disrupted staining was observed in both sALS and C9-ALS spinal cords compared to controls. Each dot represents the mean of at least two sections for each case, horizontal lines show mean and standard deviation (one-way ANOVA with Tukey *post hoc* test, n=5, 5, and 3, \**p*<0.05, \*\**p*<0.01, ns=not significant).

We also found a strong correlation between the frequency of SUN1 and SUN2 staining alteration in the spinal cords of our cohorts (r = 0.88, p < 0.0001), supporting the power of our analysis (**Supplementary Figure 7a**). Furthermore, we noted a strong correlation between Nesprin and SUN proteins disruption in each patient (**Supplementary Figure 7b-e**), which hinted at a possible causal link between the two phenotypes. In fact, and in accordance with previous literature^27^, we found that SUN1 localization at the NE is necessary for Nesprin2 protein nuclear localization. Knocking down SUN1 in HEK293 cells led to a severe depletion of Nesprin2 from the nucleus and alterations to nuclear shape (**Supplementary Figure 8**), which suggests that loss of SUN proteins from the NE may be the first pathologic change that leads to the loss of Nesprins and the disruption of LINC function.

### 4. SUN1 and SUN2 distribution is disrupted in the motor cortex of ALS postmortem tissues

As ALS patients commonly present clinically with a combination of upper and lower motor neuron signs and concomitant pathology^28^, we next performed additional immunohistochemical staining of SUN1 and SUN2 on sections from the brain motor cortex region of each case. As described above, we scored cortical neurons in the motor cortex based on the presence of a complete perinuclear ring at their equatorial plan. For both SUN1 and SUN2, we observed a significant increase in the frequency of nuclei lacking a complete perinuclear ring in sALS or C9-ALS compared with healthy controls (**Figure 5**). Furthermore, we noticed a higher degree of SUN2 staining disruption in C9-ALS compared to sALS tissues (**Figure 5h**). To further prove the strength our analyses, we looked at the relationship between the disruption of SUN1 and SUN2, finding a strong degree of correlation between these two variables (r = 0.75, *p* = 0.0030, **Supplementary Figure 9a**). We also evaluated the relationship between SUN proteins pathology across the corticospinal neural axis. For both proteins, we found a strong correlation in the degree of disruption between spinal cord (SC) and brain for each patient (SUN1: r =0.72, SUN2: r = 0.75), with the control group clearly clustering separately from both ALS cohorts (**Supplementary Figure 9 b,c**).

**Figure 5.**
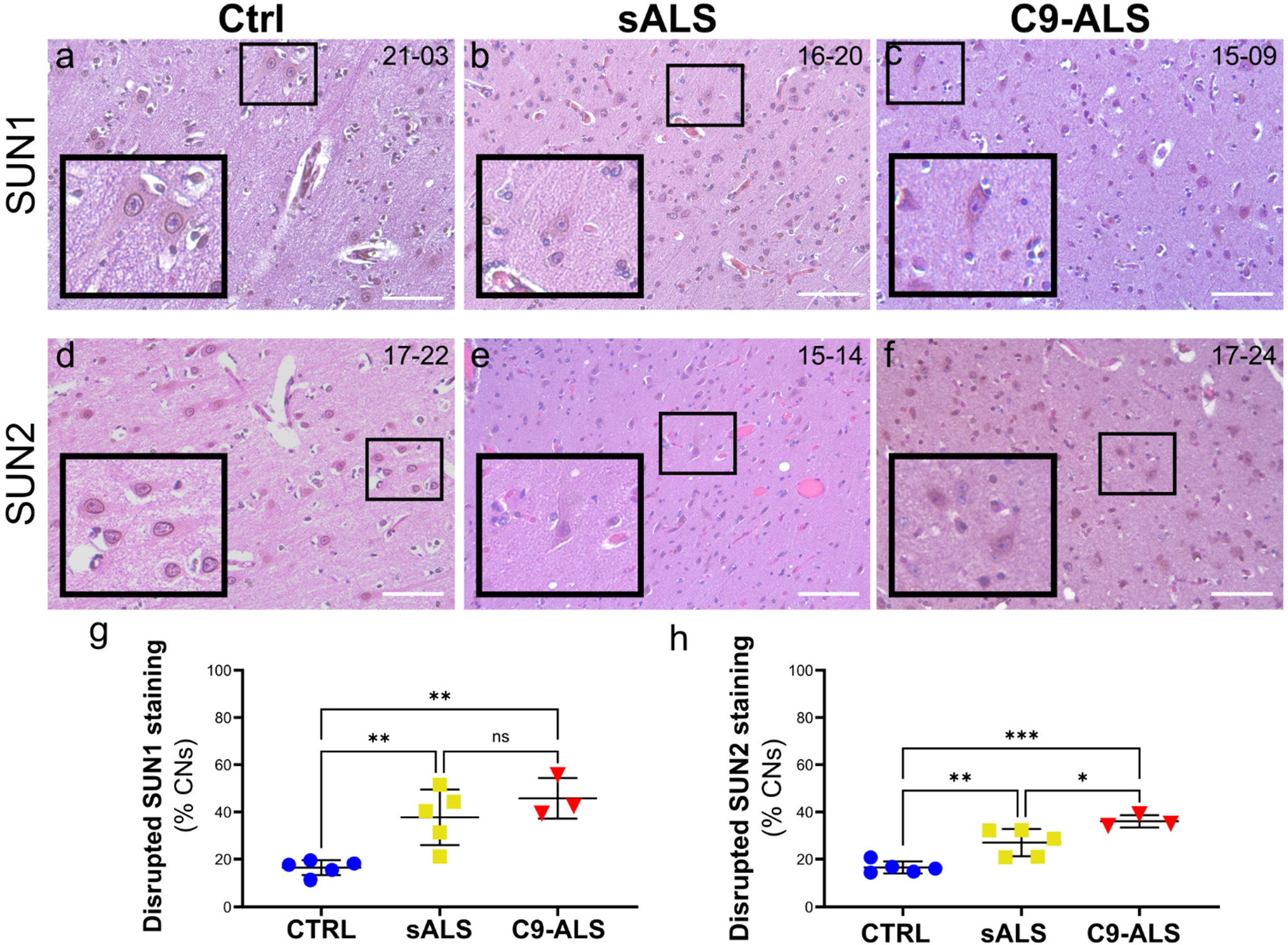
Disruption of SUN proteins in sALS and C9-ALS brain *postmortem* biopsies. **a-f**. Sections of the brain motor cortex from control (*a, d*), sALS (*b, e*) and C9-ALS (*c, f*) patients were stained with antibodies specific for SUN1 (*a-c*) and SUN2 (*d-f*). Hematoxylin and eosin counterstains were used to identify the nucleus and cytoplasm, respectively. The black boxes identify the neurons enlarged in the insets. Scale bars: 100μm. **g-h**. The frequency of disrupted NE staining for both SUN1 (*g*) and SUN2 (*h*) was quantified blindly in at least two sections from each patient’s tissue. A significant increase in the percentage of cells with disrupted staining was observed in both sALS and C9-ALS spinal cords compared to controls. A significant higher frequency of disruption in C9-ALS neurons was detected for SUN2. A similar trend was observed for SUN1, but it did not reach statistical significance. Each dot represents the mean of at least two sections for each case, horizontal lines show mean and standard deviation (one-way ANOVA with Tukey *post hoc* test, n=5, 5, and 3, \**p*<0.05, \*\**p*<0.01, \*\*\**p*<0.001, ns=not significant).

Since we found that both SUN1 and SUN2 proteins were mislocalized in cortical neurons and motor neurons *in vitro*, we wondered whether similar alterations could be found in the patients’ *postmortem* tissues. To that end, we assessed the intracellular distribution of SUN proteins by IF, using MAP2 as a cytoplasmic marker and DAPI as nuclear reference, and comparing their linear profiles between control and ALS samples (**Figure 6** and **Supplementary Figure 10**). As expected, the linear profile of both SUN1 and SUN2 in control neurons showed two high intensity peaks at the periphery of the nucleus (**Figure 6a-c** and **Supplementary Figure 10a-c**), in accordance with their localization at the NE. In sALS and C9-ALS neurons, SUN1 and SUN2 distribution showed irregular profiles lacking high intensity perinuclear peaks, confirming our blind analysis of the immunohistochemistry data (**Figure 6d-i** and **Supplementary Figure 10d-i**). Furthermore, a greater frequency of high-intensity peaks was noted outside of the nucleus. Together, these data demonstrate a severe impairment in the localization of SUN proteins to the NE in the motor cortex of ALS patients, leading to mislocalization and accumulation of both proteins in the cytoplasm of these cortical neurons.

**Figure 6.**
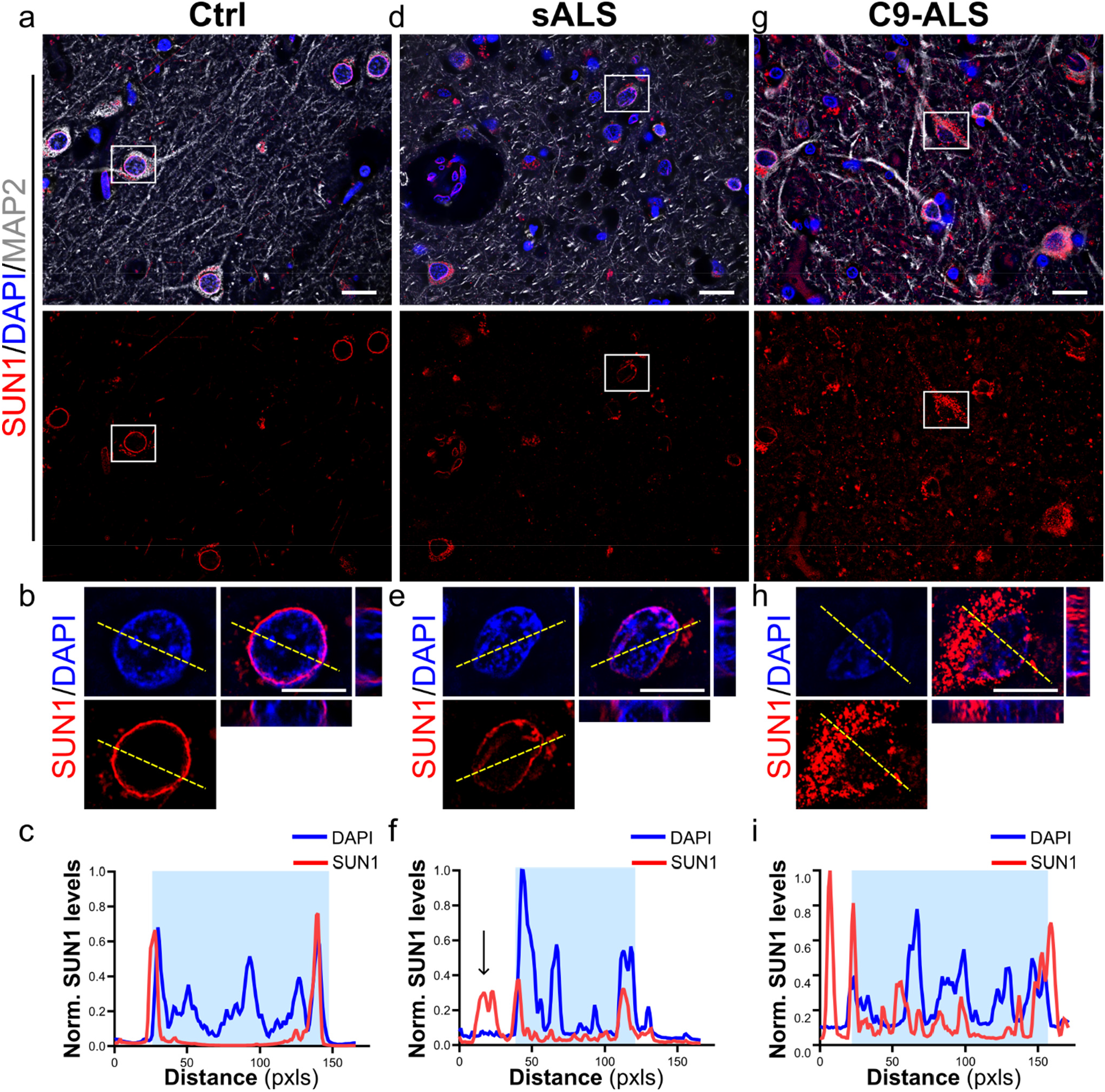
SUN1 is mislocalized in the cytoplasm in cortical neurons of sporadic and C9-ALS patients. Representative images of motor cortex sections from control (***a, b***), sALS (***d, e***) and C9-ALS (***g, h***) patients stained with antibodies specific for SUN1 (*red*) and MAP2 (*grays*). DAPI (*blue*) was used to label nuclei. White boxes in *a, d*, and *g* identify neurons enlarged in *b, e*, and *h*. Line profile plots show the marked difference of SUN1 distribution in sALS (***f***) and C9-ALS (***i***) cortical neurons compared to controls (***c***). The yellow dashed lines in *a* indicate the lines used for the profile plots. Scale bars: 20μm in main panels, 10μm in zoomed-in images.

### 5. LINC complex disruption contributes to altered nuclear morphology in ALS neurons

Since the LINC complex is a known regulator of nuclear morphology and size through its interaction with the nuclear lamina and cytoskeleton^29^, we wondered whether the observed alterations in the levels and localization of its main components could impact the control of nuclear size. To test that possibility, we first quantified the area of the nucleolus at its equatorial plane in our 2D and 3D cultures and normalized it to the area of the whole nucleus (relative nucleolar area). While we did not observe any major alteration in iPSC-derived MNs grown on a solid 2D substrate (not shown), we found that ISL1-positive C9 MNs grown in the organoids had significantly smaller nucleoli compared to their isogenic controls (**Supplementary Figure 11a-c**), in agreement with previously reported *in vivo* observations. When we performed a similar analysis in the spinal cord biopsies, we again observed in both sALS and C9-ALS motor neurons a significant reduction in the absolute nucleolar area (13.99 ± 4.75 μm for sALS, 9.73 ± 3.93 μm for C9-ALS and 22.71 ± 4.77 μm for CTRL) and a trend toward smaller nuclei (166.8 ± 40.80 μm for sALS, 156 ± 51.36 μm for C9-ALS and 204.5 ± 40.61 μm for CTRL) compared to non-neurological controls (**Supplementary Figure 11d-f**). No difference was observed in the average cytoplasmic area (879.7 ± 183.2 μm for sALS, 826.6 ± 373.8 μm for C9-ALS and 881.4 ± 167.3 μm for CTRL).

To find out a possible causal relationship between LINC complex disruption and nuclear morphological features, we compared nuclear and nucleolar areas between ALS neurons with normal and disrupted SUN staining. For this analysis, we normalized the nuclear area to the area of the cytoplasm, and the nucleolar area to the area of the nucleus, to avoid confoundings due to differences in cell body or nuclear size (**Figure 7**). Interestingly, we found that both sALS and C9-ALS neurons with disrupted SUN1 or SUN2 staining had significantly smaller nucleus to soma ratios compared to control neurons (**Figure 7b, d**). An even bigger difference was found when the normalized nucleolar area was compared between control and SUN1 or SUN2-disrupted ALS MNs (**Figure 7d, e**). Importantly, we found that sALS or C9-ALS MNs with intact SUN1 or SUN2 staining had instead comparable nuclear and nucleolar areas to the healthy controls (**Figure 7b-e**), suggesting that LINC complex disruption may be driving severe alterations to nuclear and nucleolar homeostasis.

**Figure 7.**
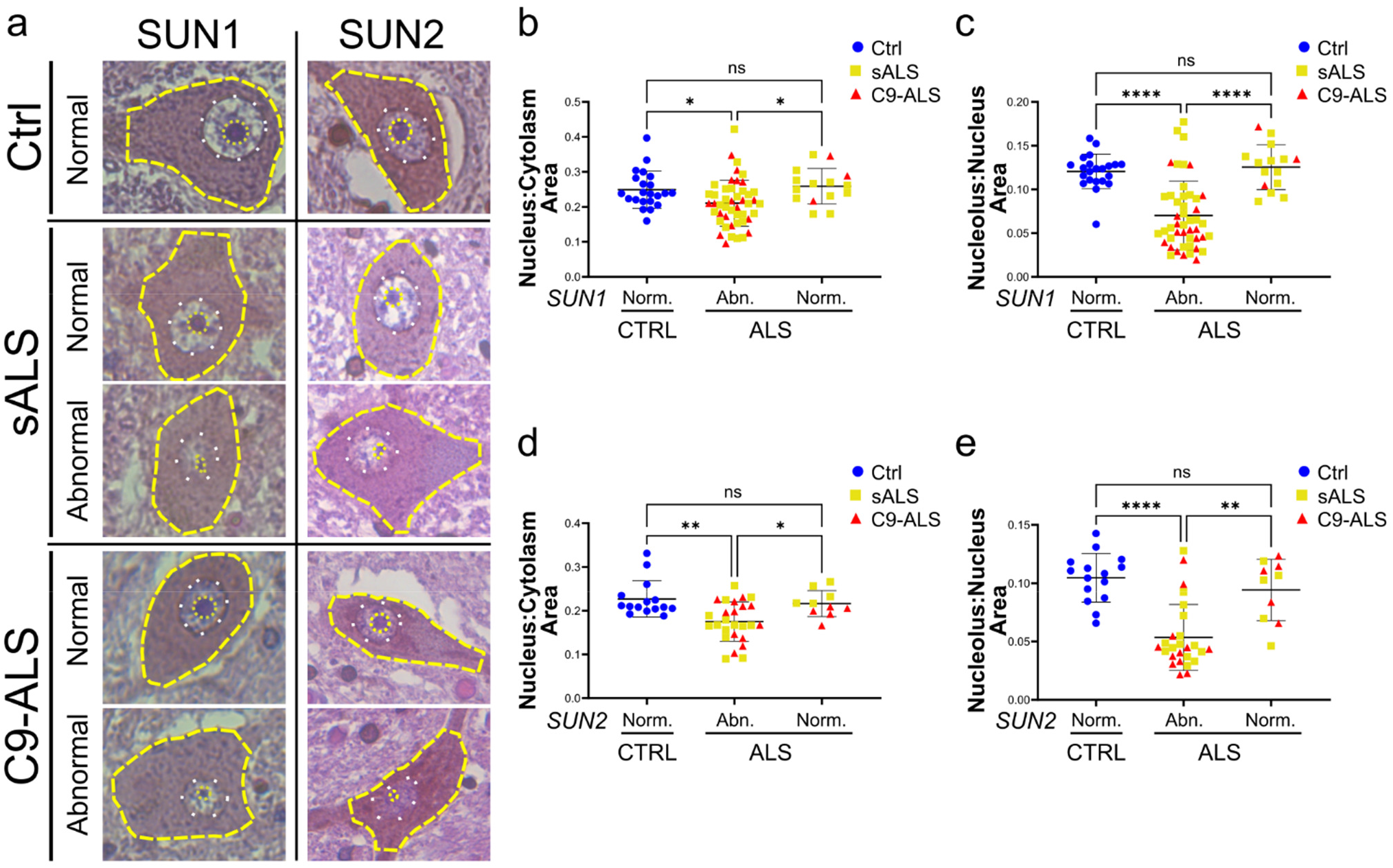
LINC complex disruption correlates with nuclear morphological alterations in ALS MNs. **a**. Representative images of spinal motor neurons from control (*top*), sALS (*middle*), and C9-ALS (*bottom*) with normal or abnormal SUN1 or SUN2 staining. The dashed yellow lines indicate the cells contour, white dots identify the nucleus, and yellow dots highlight the nucleolus. **b-e**. The quantification of the relative nuclear (*b, d*) and nucleolar (*c, e*) area in cells categorized based on the presence of a normal or abnormal SUN1 (*b, c*) or SUN2 (*c, e*) nuclear staining shows a significant correlation between LINC complex disruption and smaller nuclear and sub-nuclear structures (one-way ANOVA with Tukey *post hoc* test, n=22, 47, and 15 in *b* and *c*, n=15, 26, and 10 in *d* and *e*, \**p*<0.05, \*\**p*<0.01, \*\*\*\**p*<0.0001, ns= not significant).

We next wondered what role, if any, could TDP-43 nuclear depletion and aggregation, the main pathological hallmark of ALS/FTD, have in driving LINC complex disruption and the LINC-associated decrease in nuclear and nucleolar size. To address this important question, we focused our analysis on the motor cortex, since we observed a broader range of LINC complex pathology in this tissue compared to the spinal cord, where most MNs presented with severe alterations of all LINC proteins. When we quantified the co-occurrence of TDP-43 nuclear depletion and SUN1 disruption in MAP2-positive cortical neurons, we found that almost the totality of the ALS neurons with mislocalized TDP-43 also presented with SUN1 perinuclear disruption (**Supplementary Figure 12**), supporting a strong correlation between the two events. However, we also found that about 20% of neurons with normally localized TDP-43 presented alterations in SUN1 distribution (**Supplementary Figure 12**), suggesting that either LINC complex alterations precede TDP-43 mislocalization, or that the two phenomena are independent from each other.

Based on these results, we decided to evaluate changes to the nuclear morphology (i.e., area and circularity) and soma size in cortical neuron divided into four sub-groups: (1) Control neurons, used as a reference, (2) ALS neurons with normal SUN1 and TDP-43 staining, (3) ALS neurons with disrupted SUN1 staining but normal TDP-43 distribution, and (4) ALS neurons with disrupted SUN1 staining and abnormal cytoplasmic TDP43 mislocalization. Similar to what we observed in spinal MNs, we found that ALS neurons with normal SUN1 distribution displayed nuclear parameters indistinguishable from non-neurological controls (**Figure 8**). Disruption of SUN1 staining instead correlated strongly with a significant reduction in nuclear circularity and area, which was irrespective of changes in TDP-43 localization (**Figure 8c and Supplementary Figure 13b**). However, our analyses also revealed that TDP-43 cytoplasmic mislocalization not only further reduced absolute nuclear size (**Figure 8c**), but also led to a significant reduction in the overall size of the cell soma (**Supplementary Figure 13a**), suggesting that TDP-43 nuclear depletion and cytoplasmic aggregation may lead to general cell toxicity which impacts many different aspects of cellular homeostasis. This observation also explains the paradoxical trend of the relative nuclear size in ALS neurons, which is heavily reduced in neurons with disrupted SUN1 but normal TDP-43 staining but seems to return to control levels in neurons with cytoplasmic mislocalization of TDP-43 (**Figure 8b**). Overall, our results suggest that LINC complex disruption may be an early event that leads to the pathological alteration of nuclear homeostasis, contributing to ALS/FTD pathogenesis.

**Figure 8.**
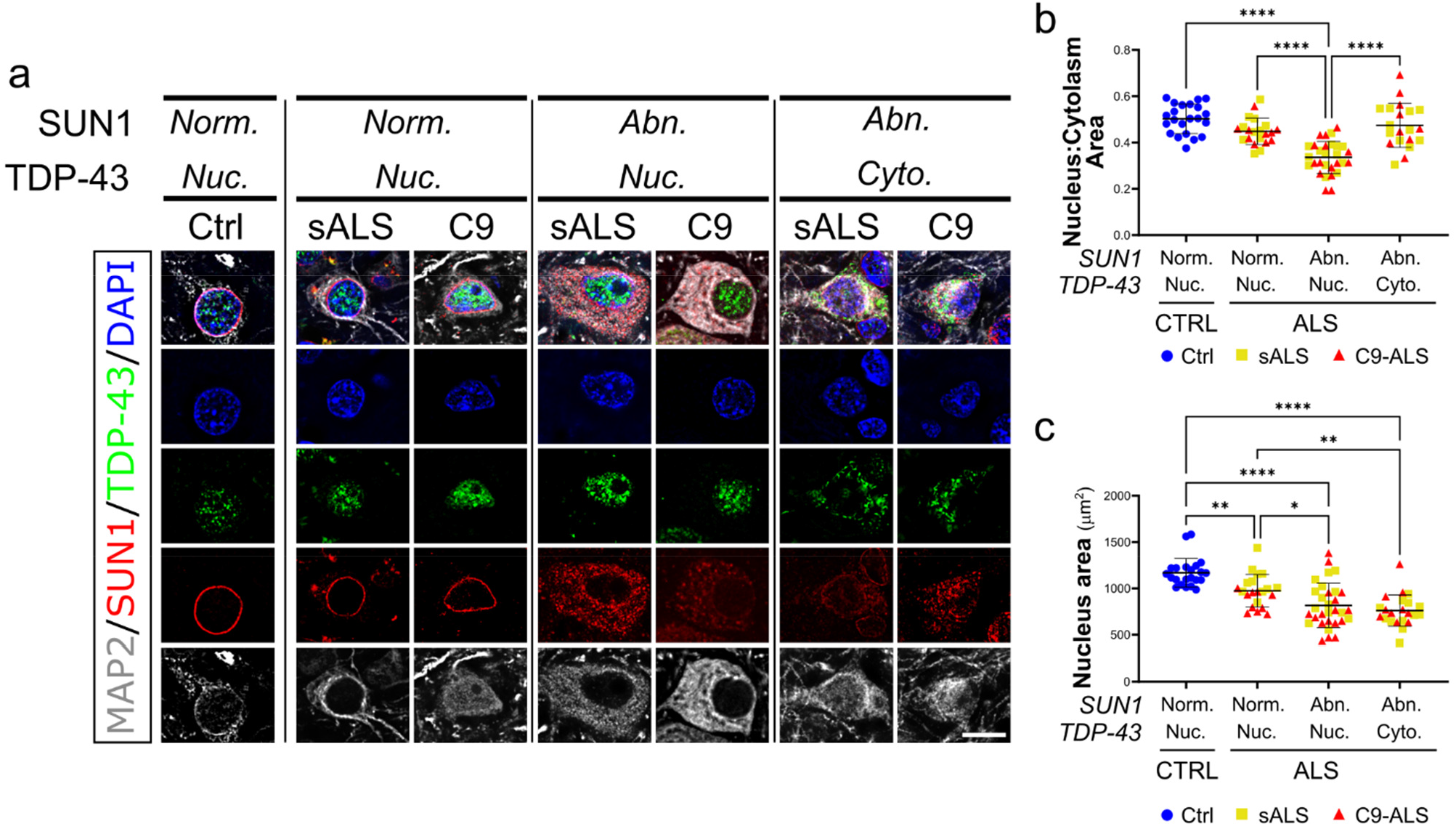
SUN1 nuclear loss is a main contributor to cellular morphological alterations. **a**. Representative images of cortical neurons categorized based on the presence of nuclear (Nuc.) or cytoplasmic (cyto.) TDP-43 (*green*) and normal (Norm.) or abnormal (Abn.) SUN1 staining (*red*). MAP2 (*grays*) was used as a neuronal marker, while DAPI (*blue*) labeled the cell nucleus. Scale bar: 10μm. **b**. Quantification of the relative nuclear to cytoplasmic area shows that neurons with abnormal SUN1 distribution have significantly smaller nuclei compared to all other groups (one-way ANOVA with Tuckey post *hoc* test, n=22, 21,29, 20, \*\*\*\**p*<0.0001). **c**. Quantification of the absolute nuclear area of cortical neurons shows that all ALS neurons have smaller nuclei compared to healthy controls. However, loss of SUN1 from the NE (categorized as abnormal) further impacts nuclear size, regardless of TDP-43 localization (one-way ANOVA with Tuckey post *hoc* test, n=22, 21,31, 22, \**p*<0.05, \*\**p*<0.01, \*\*\*\**p*<0.0001).

## Discussion

Recent studies in several models of sporadic and familial ALS and FTD, including mutant *C9ORF72, TDP-43, FUS* and *PFN1*, have pointed to the loss of nuclear envelope (NE) and NPC integrity, as well as disruption of nucleocytoplasmic transport, as key drivers of disease initiation and progression^23,30,31,32^. While many of the molecular players and pathways that lead to such dysfunctions are currently unknown, our previous work has pointed to the nucleus-cytoskeleton connection as a possible mediator of this process. In fact, we have previously shown that positive or negative modulation of actin polymerization can lead to NPC injury and nuclear import defects^32^. Furthermore, we recently showed that increased mechanical tension on the cell’s nucleus can lead to breaks in the NE, NPC injury, and DNA damage^6^. The LINC complex is a structural component of the nuclear envelope and a fundamental element in the mechanical transmission of physical stimuli to the nucleus. Through its interactions with the cellular cytoskeleton, nuclear lamina, and NPC, the LINC complex plays a pivotal role in the maintenance of nuclear homeostasis in response to mechanical strain^33^. Together, these observations prompted us to directly investigate the role of the LINC complex in ALS/FTD disease.

In this study, we interrogated three different ALS/FTD models, including iPSC-derived cortical and motor neurons, spinal cord organoids, and *postmortem* sALS and C9-ALS patient-derived brain and spinal cord biopsies, and identified severe alterations in the nuclear levels and cellular distribution of the four most abundant components of the LINC complex: SUN1, SUN2, Nesprin1, and Nesprin2. Our analyses of *postmortem* tissues provided important insights into the relevance of LINC complex disruption to human disease. However, patient-derived samples often represent a late stage of the disease with extensive degeneration, which may not exhibit molecular or cellular signatures directly associated with the initiating events that caused the disease. By contrast, neurons derived from iPSCs, while immature and lacking normal tissue connections, can provide insights into the earliest stages of neurodegeneration, shedding light on the pathogenic events that drive disease initiation. By combining both approaches, our data rigorously show that cellular and/or NE levels for all four LINC proteins tested were severely reduced both in cortical and motor neurons. A similar reduction in SUN1 levels have been recently reported in iPSC-derived spinal neurons^19^, which confirms and strengthens our observations. We also found that spinal motor neurons in the *postmortem* tissues displayed a higher degree of disruption compared to cortical neurons, but the levels of LINC complex pathology between spinal cord and brain from each individual patient correlated strongly, indicating a parallel evolution, albeit of different magnitude, of this phenotype in different compartments of the CNS affected by ALS/FTD. Our comprehensive approach also allowed us to investigate the relationship between the inner nuclear membrane SUN proteins and the outer nuclear membrane Nesprin proteins. Interestingly, we found a strong correlation between SUN and Nesprin proteins’ alteration across all models, suggesting that SUN protein localization at the NE is necessary for Nesprins assembly in the LINC complex^27^, and that loss of SUN proteins may trigger Nesprin mislocalization and loss of LINC complex activity.

Through its interaction with the cytoskeleton and nuclear lamina, the LINC complex plays a fundamental role not only in the modulation of nuclear positioning, but also in regulating nuclear morphology and in promoting changes and adaptations to chromatin organization^29^. For instance, acute depletion of SUN1 was shown to alter nuclear morphology, chromatin organization, and nucleolar distribution^11^, while deletion or overexpression of different LINC proteins was proven to alter the ratio between nuclear and cytoplasmic volumes^34^. It has been recently demonstrated that nuclear and nucleolar shrinkage occurs in both C9-ALS and sALS spinal MNs independently of many other common pathological changes, including TDP-43 mislocalization or HRE RNA foci^35^. Nucleolar size alterations have also been observed in C9-ALS patient lymphocytes, fibroblasts and patient iPSC-derived neurons^36^. Surprisingly, in these cellular models the nucleoli were found to be larger when compared to controls, suggesting that either aging or the non-physiological conditions of 2D cell culture may affect the phenotype differently. However, the cause of this phenomenon is not well understood. Our analyses of the impact of LINC complex alterations on nuclear morphology showed a strong correlation between SUN1/SUN2 disruption and the reduction of nucleolar and nuclear size in ALS MNs. While MNs with SUN1 and SUN2 normally localized at the NE displayed nuclear and nucleolar sizes comparable to the MNs of non-neurological controls, altered SUN1/SUN2 staining strongly correlated with smaller nucleoli and an overall reduced nuclear volume, suggesting that alterations to the LINC complex may be driving this phenotype. Interestingly, we were able to observe nuclear and nucleolar alterations both in patient-derived tissues and 3D spinal organoids, while 2D-cultured motor or cortical neurons lacked a well-defined and mature nucleolus and did not display obvious alterations to nuclear morphology under our experimental conditions. While 2D cultures are a valuable tool that allow for the isolation of cell-autonomous events by limiting confounding variables, they nonetheless represent a reductionist model that lacks the complex interactions of cells with their environment, including cell-to-cell and cell-extracellular matrix (ECM) mechanical interaction. For our study in particular, such interactions were particularly relevant, given the well-established role of the LINC complex in transmitting forces from the ECM to the nucleus to modulate cell mechanical properties and function. Spinal cord organoids thus offered a more complex model that could better recapitulate both tissue cell diversity and complex mechanical interactions between cells and their surroundings ^37^.

One important and surprising observation was that most of the LINC-dependent phenotypes we observed, including protein mislocalization and nuclear morphology alterations, were independent from any TDP-43 related pathology. While rare co-aggregation of SUN1 or SUN2 with TDP-43 was observed, the majority of mislocalized cytoplasmic SUN proteins formed discrete clumps that did not contain TDP-43, and most TDP-43 skein-like aggregates were negative for SUN proteins. Similarly, nuclear and nucleolar shrinkage did not correlate with TDP-43 nuclear depletion or aggregation, which instead caused an overall reduction in cellular volume. Shrinkage of nuclear and nucleolar size has been previously described in ALS *postmortem* tissue^35,38^ and prp-TDP-43^A315T^ mouse model, seemingly preceding neuronal loss and motor defects^38^. Our novel data provide a mechanistic explanation for this early sign of disease and suggest that LINC complex defects may be an early event in the pathogenic cascade in ALS/FTD, representing a parallel pathological event that adds to the burden of disease caused by TDP-43-related dysfunction.

In conclusion, our results, obtained in complementary *in vivo* and *in vitro* models, provide a robust body of evidence pointing at the alteration of the LINC complex as an early ALS phenotype conserved in the late stages of the disease, making the LINC complex a possible target for both biomarker or therapy development.

## Methods

### iPSCs culture

The protocols met all ethical and safety standards for work on human cells. The human iPSC lines used in this study are identified in **Supplementary Table 1** and were fully characterized in our lab. They were maintained on Matrigel-coated plates in StemFlex Medium (Gibco), and passaged every 4–6 days using 0.5mM EDTA in Ca^2+^ and Mg^2+^-free 1x PBS (Thermo Fisher Scientific).

### Cortical neuron (i^3^CN) differentiation

To differentiate iPSCs to i^3^CNs, we used the method described by Dr. Michael E. Ward^25^. This method relies on the integration of a gene expression cassette in the safe-harbor CLYBL1 locus and produces >90% pure neuronal populations. To integrate the cassette, two independent pairs of isogenic *C9ORF72* iPSCs were transfected with a plasmid containing the coding sequence for the NGN2 transcription factor, NLS-mApple fluorescent protein, and blasticidin resistance gene, flanked by homology arms to the CLYB1 genomic locus (Addgene plasmid # 124229), and ribonucleoprotein particles containing the Cas9 nuclease and a site-specific gRNA (Synthego). Colonies were positively selected based on blasticidin (10μg/ml) resistance and the expression of NLS-mApple. To minimize the effects of inter-clonal variation, all cells positive for the cassette integration were pooled together and expanded as neuroprogenitors. To induce differentiation, 2 μg/ml doxycycline was added to the medium for 2 days, while proliferating cells were killed off with BrdU treatment. Cells were than plated on poly-ornithine and poly-lysine coated coverslips and switched to neuronal medium (Neurobasal, 2% B27, 1% N2, 1% NEAA, 1μg/ml laminin). To verify full differentiation, positivity for NeuN and Tuj1 and negativity for Oct4 and Sox2 was assessed.

### Motor neuron (iMN) differentiation

iMNs were generated following the protocol published by Du et al.^39^ with minimal modification. Briefly, confluent iPSC cultures were detached using 0.5mM EDTA and plated 1:4 on Matrigel ES-coated plates in StemFlex Medium (Gibco) supplemented with 10μM ROCK inhibitor (SellChem). The following day, cells were switched to neural induction media (DMEM/F12 and Neurobasal medium at 1:1 ratio, 1% B27, 0.5% N2, 1% PenStrep, 1% Glutamax; ThermoFisher Scientific) supplemented with ascorbic acid (0.1 mM, Sigma-Aldrich), CHIR99021 (3 μM, Tocris), DMH1 (2 μM, Tocris) and SB431542 (2 μM, Tocris). On day 7 and 13, cells were dissociated with Accutase (Millipore) and plated on Matrigel growth factor reduced (MaGR, Corining) in neural induction media further supplemented with 0.1 μM retinoic acid (Sigma-Aldrich) and 1 μM purmorphamine (Calbiochem). Valproic acid (0.5mM, Tocris) was added on day 13. To induce MN differentiation, cells were dissociated with Accutase (Millipore) and cultured in suspension for 6 days on an horizontal shaker in neural medium supplemented with 0.5 μM retinoic acid and 0.5 μM purmorphamine. On day 25, cells were dissociated into single cell suspension with Accutase (Millipore) and plated on MaGR-coated plates in neural medium with 0.5 μM retinoic acid, 0.5 μM purmorphamine and 0.1 μM Compound E (Calbiochem) until day 35.

### Spinal cord organoids

Spinal cord organoids were generated following published protocols with minor modifications ^26^. We first dissociated iPSCs into single cells, seeded 30,000 cells per well in Nunclon Sphera ultralow attachment 96-well plates (ThermoFisher Scientific) in neural induction medium supplemented with 10 ng/mL FGF (Corning), 20 ng/mL EGF (Corning) and 10 μM ROCK inhibitor. Neuralization was induced via treatment with 3 μM CHIR99021, 2 μM DMH1 and 2 μM SB431542. Media was changed every other day and supplementaed with 0.1 μM retinoic acid from day 3 to induce caudalization. At day 10, organoids were embedded in 1% Matrigel Matrix for Organoid Culture (Corning) and kept in slow (70rpm) horizontal agitation. From day 10, neural media was further supplemented 0.5 μM retinoic acid and 1 μM purmorphamine, while 0.1 mM ascorbic acid, 20 ng/mL BDNF, 10 ng/mL GDNF were added after day 20 and till day 120.

### HEK293 cell culture and transfection

HEK293 cells were grown in DMEM media supplemented with 10% FBS. For immunofluorescence experiments, cells were plated on poly-lysine coated coverslips (1mg/ml; Millipore Sigma) and allowed to recover for 24 hours. Cell transfection was carried out using TurboFect reagent (ThermoFisher Scientific) according to manufacturer’s instructions using following plasmids: pLV[shRNA]-EGFP-U6>hSUN1 (target sequence: TTCATGGACGAGGGCATATAC) or pLV[shRNA]-EGFP/Puro-U6>Scramble_shRNA (VectorBuilder). Cells were processed for immunofluorescence 48 hours after transfection.

### Immunofluorescence and image acquisition of in vitro models

Motor or cortical iPSC-derived neurons were seeded on Matrigel (Corning) or poly-D-Lysine (Millipore Sigma) coated glass coverslips at the density of 300 cell/mm^2^. Cells were fixed with 4% paraformaldehyde for 15 min and permeabilized with 0.2% Triton-X 100 for 15 min. Cells were blocked with 5% bovine serum albumin before hybridization with the appropriate antibodies overnight at 4 °C. Anti-mouse and anti-rabbit donkey secondary antibodies conjugated with either Alexa Fluor 647, Rhodamine-Red X, or Alexa Fluor 488 (Jackson Immunoresearch and ThermoFisher Scientific) were hybridized for 1 h at room temperature.

Organoids were washed in PBS, fixed in 4% PFA at 4°C overnight, and incubated in 30% sucrose for cryopreservation for at least 16 hours. Organoids were embedded into OCT and preserved at -80°C. Sections were sliced at 10 μm thickness using a cryostat (Leica), permeabilized with 0.1% Triton-X 100 for 10 minutes and incubated in blocking buffer (5% FBS, 1% BSA, 0.01% Triton-X 100) for 2 hours. Slides were incubated with primary antibodies (see **Supplementary Table 4**) overnight at 4°C, followed by 2 hours hybridization with appropriate secondary antibodies at room temperature.

Organoids sections and cells coverslips were mounted onto a glass slide using Prolong Gold mounting medium (ThermoFisher Scientific) and imaged using a widefield microscope (Leica DMi8 Thunder) equipped with a cooled CMOS camera (DFC9000 GTC). Images were acquired as Z-stacks (0.21 μm step size) using a x63 lens unless otherwise specified.

### Image analysis

Immunofluorescence images were deconvolved using an adaptative blind deconvolution algorithm (Autoquant X3, Media Cybernetics) before analysis. Fluorescence intensity levels were quantified using Fiji^40^ (US National Institutes of Health). First, 3D stacks were compressed to 2D images using the maximal intensity projection algorithm. Second, a mean filter (with radius 2.0 pixels) was applied to the DAPI nuclear staining channel to discretely separate single nuclei. Finally, mean fluoresce levels (MFI) of the target proteins were measured within discrete region of interests (ROI) established by gating single nuclei.

For the analysis of linear profiles of SUN and Nesprin proteins, the equatorial plane of the cells was manually identified, and a horizontal line was drawn across the center of the DAPI nuclear staining and through the entire cell. Linear profiles of DAPI and LINC complex proteins staining were generated using Fiji Plot Profile function, normalized to controls, and then merged on the same graph to be compared.

To evaluate qualitative alterations in the nuclear distribution of LINC proteins, a blinded analysis was performed on 3D stacks of individual optical slices. To avoid misinterpretation of staining profiles, no 3D to 2D compression of the images was performed as suggested^41,42^. For LINC complex analysis, abnormal staining was considered if the signal was not uniformly distributed around the nucleus with the presence of gaps. For all experiments, raw values were normalized to the mean of the control condition.

### Western Blots

Cells were centrifuged at 250 x g for 10 min at 4°C and processed in lysis buffer (10 mM Tris-HCL pH 8, 100 mM NaCl, 1 mM EDTA pH 8, 1% NP40) supplemented with protease inhibitor (Complete EDTA-free, Roche) and phosphatase inhibitor (phosphoSTOP, Roche) and sonicated (40 kHz for 10s; Sonifier® SFX150, Emerson) to ensure uniform and complete lysis. Protein extracts (30 μg) were resolved by SDS-PAGE on Mini 4% - 20% Novex Tris-Glycine Gels (Thermo Fisher). Primary antibodies were incubated overnight at 4°C (**Supplementary Table 4**). Secondary antibodies conjugated with IRDye® infrared fluorophores (LI-COR) were incubated for 1 h at room temperature. Blots were visualized using the Odyssey Infrared Imaging System (LI-COR).

### Immunohistochemistry and immunofluorescence of Human postmortem samples

Human postmortem motor cortex and cervical spinal cord paraffin sections were obtained from TargetALS biorepository. Clinical data from the cases are summarized in **Supplementary Table 2** and **Supplementary Table 3**. Sections were de-paraffinized in xylene (Sigma-Aldrich) and re-hydrated with scaling dilutions of ethanol (ThermoFisher Scientific) to MilliQ water. Endogenous peroxidase blocking was conducted with 0.3% H_2_O_2_ in Methanol solution for 40 minutes (ThermoFisher Scientific) followed by re-hydration with scaling concentrations of methanol. Antigen retrieval was performed with 10 mM citric acid pH 6 at 100 °C for 15 minutes. After an hour of cool-down slides were washed with PBS and incubated with blocking buffer (1% BSA, 5% FBS, 0.01 % Triton-X 100 in 1x PBS) followed by Avidin/Biotin blocking (Vector laboratories). Slides were incubated overnight at 4°C with primary antibodies (see **Supplementary Table 4**) and at room temperature for 1 hour with secondary biotinylated antibody (anti-rabbit IgG (H+L), Jackson Laboratories). For immunoreactivity detection, slides were incubated with Vectastain Ekite ABC reagent (Vector laboratories) for 30 minutes and then with ImmPACT DAB substrate kit (Vector laboratories) from 2 to 5 minutes. After counterstain with hematoxylin and/or eosin, slides were de-hydrated and mounted on microscope cover glass (Globe Scientific) with mounting medium (Epredia). Sections were imaged using a widefield microscope (Leica DMi8 Thunder) equipped with a color digital camera (DMC5400) and a 20x objective. For quantification of nuclear, nucleolar, and somatic areas, hematoxylin was used as nuclear marker while eosin was used to identify the whole cytoplasm. For immunofluorescence assays of *postmortem* tissue sections, we followed the same protocol already described for organoids sections after the initial steps of de-paraffinization and de-hydration.

## Acknowledgements

This work was supported by the National Institute of Neurodegeneration and Stroke (NINDS) within the National Institute of Health (NIH) under Grant # R01NS116143 and the RI-INBRE Early Career Development Award to CF. Access to core facility was allowed through the Institutional Development Award (IDeA) Network for Biomedical Research Excellence from the National Institute of General Medical Sciences (NIGMS) of the NIH under grant # P20GM103430. We are indebted to Target ALS and ALS patients for postmortem tissue donation, and to Dr. Ward (NIH) and Dr. Talbot (University of Oxford, UK) for sharing the *C9ORF72* mutant and isogenic control iPSC lines. The CLYBL-TO-hNGN2-BSD-mApple was a gift from Michael Ward (Addgene plasmid # 124229). The authors have no relevant financial or non-financial interests to disclose.

