## Supplementary material for "LINC complex alterations are a hallmark of sporadic and familial ALS/FTD"

**Content:**


- **Supplementary Figure 1.** iPSC-derived motor neuron differentiation.
- **Supplementary Figure 2.** SUN1 and Nesprin2 disruption in iPSC-derived iMNs.
- **Supplementary Figure 3.** iPSC-derived cortical neuron differentiation.
- **Supplementary Figure 4.** SUN2 and Nesprin1 alterations in C9-ALS $i^3$CNs.
- **Supplementary Figure 5.** iPSC-derived spinal cord organoids differentiation.
- **Supplementary Figure 6.** SUN2 and Nesprin2 disruption in sALS and C9-ALS spinal cord *postmortem* biopsies.
- **Supplementary Figure 7.** Correlative analysis of LINC protein disruption in spinal cord biopsies.
- **Supplementary Figure 8.** SUN1 is fundamental for Nesprin2 nuclear envelope localization.
- **Supplementary Figure 9.** Correlative analysis of SUN proteins disruption.
- **Supplementary Figure 10.** SUN2 mislocalization in the cytoplasm in cortical neurons of sporadic and C9-ALS patients.
- **Supplementary Figure 11.** Loss of SUN1 correlates with altered nuclear and nucleolar size.
- **Supplementary Figure 12.** TDP43 aggregation strongly associates with SUN1 disruption.
- **Supplementary Figure 13.** Morphological changes in cortical neurons depend on both TDP-43 and SUN1.

- **Supplementary Table 1.** List of iPSC lines used in the study.
- **Supplementary Table 2.** Clinical and pathological information of non-neurological control cases.
- **Supplementary Table 3.** Clinical and Pathological information of ALS cases.
- **Supplementary Table 4.** Detailed information on reagents and resources used.

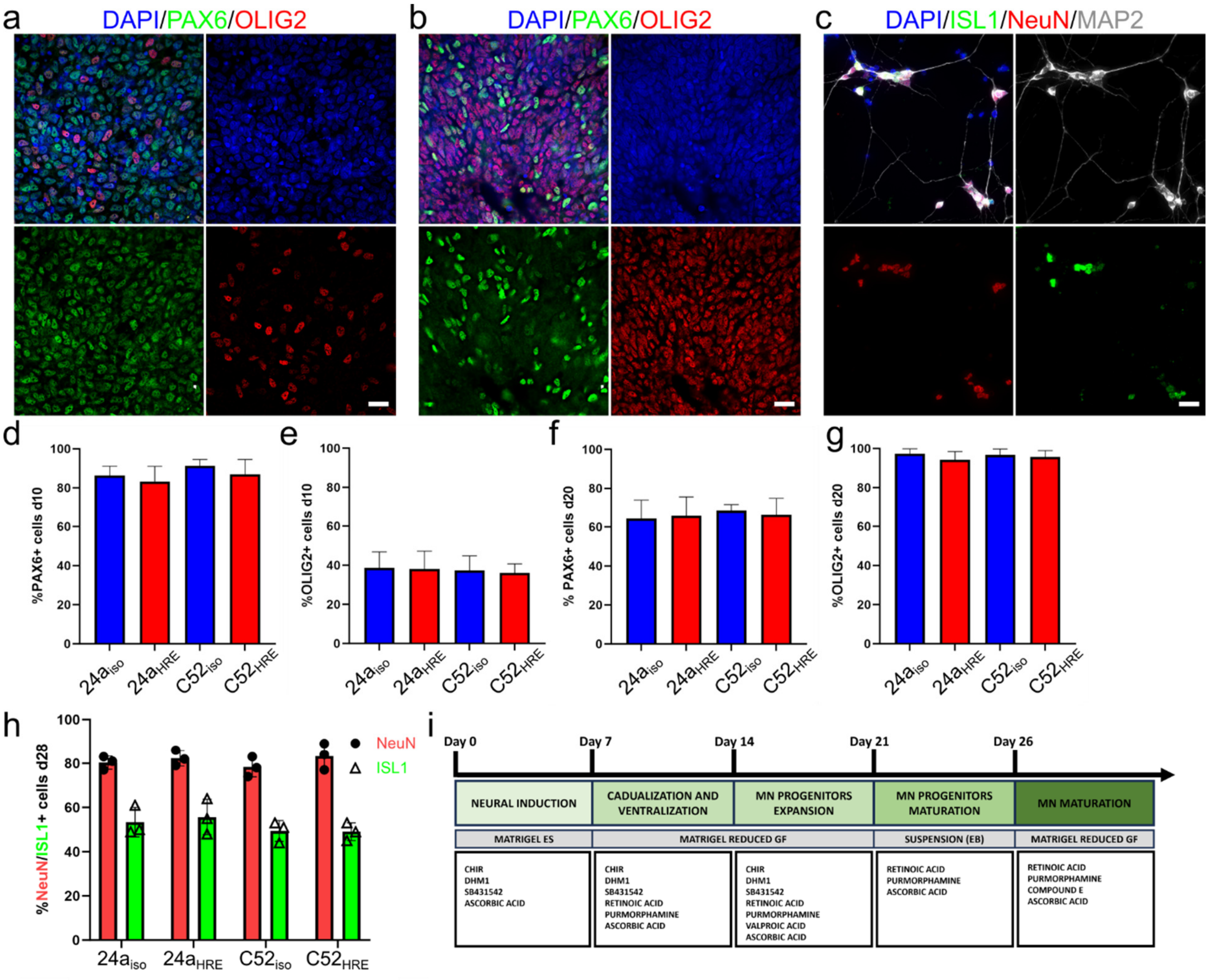


**Supplementary Figure 1. iPSC-derived motor neuron differentiation. a-c** Representative images of iPSC-derived motor neurons (iMNs) 10 days (*a*), 20 days (*b*) and 28 days (*c*) after the induction of differentiation. Scale bars: 20 µm. **d-e.** At day 10, neuroprogenitors cells (NPCs) display a strong immunoreactivity for PAX6 (~85%, green in *a*), while a small fraction of cells expresses OLIG2 (~35%, red in *a*). **f-g.** At day 20, PAX6 immunoreactivity is decreased (~60%, green in *b*) while almost the totality of the cells is OLIG2-positive (~95%, red in *b*), indicating that cells have entered the maturation phase. **h.** At day 28, immature iMNs start to be positive for the neuronal marker NeuN (~85%, red in *c*) and the MN-specific marker ISL1 (~55%, green in *c*). For all markers, no significant difference in the efficiency of differentiation was observed among the different iPSC lines. **i.** Schematic representation of the iMN differentiation protocol.

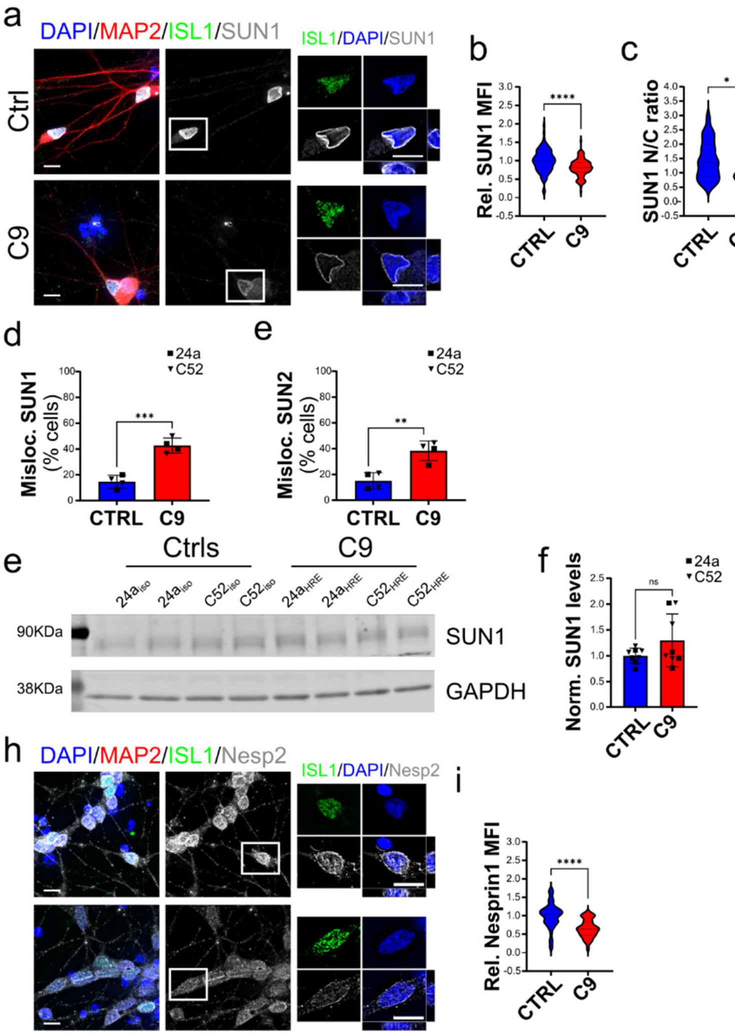


**Supplementary Figure 2. SUN1 and Nesprin2 disruption in iPSC-derived iMNs. a.** Representative images of SUN1 staining (*grays*) in control and C9 iMNs. **b-d.** Quantification of SUN1 levels shows significant downregulation in its nuclear mean fluorescence intensity (MFI, *b*), a decrease in nucleus to cytoplasm (N/C) ratio (*c*), and an increase in the frequency of SUN1 mislocalization from the NE (*d*) mutant C9 cells compared to isogenic controls. Data were aggregated from 4 independent experiments in both 24a and C52 isogenic pairs. Student's *t* test; n=94 and 102 in *b*, n=50 and 47 in *c*, n=4 in *d*; **p*<0.05, ****p*<0.001, *****p*<0.0001. **e.** The frequency of SUN2 mislocalization from the NE in mutant cells compared to isogenic controls was similarly increased. Student's *t* test; n=4, ***p*<0.01. **f-g.** Representative blot and quantification of the levels of SUN1 in whole cell lysates show no significant change in overall levels. Data were collected from 5 independent experiments from both isogenic pairs. Student's *t* test; n=8; ns, not significant. **h-i.** Representative images (*h*) and quantification (*i*) of Nesprin2 staining in iMNs from C9 mutant and isogenic lines show significant reduction in Nesprin2 nuclear MFI. For both *a* and *h*, Islet1 (ISL1, *green*) and MAP2 (*red*) were used to identify motor neurons. DAPI (*blue*) labeled the nucleus. The white boxes indicate the neuron enlarged in the panels on the right. Student's *t* test; n=65 and 43 from 4 independent experiments; *****p*<0.0001. For all, bars represent mean and SEM, violin plots show data distribution with dashed lines indicating median and quartile range. Scale bars are 10 μm.

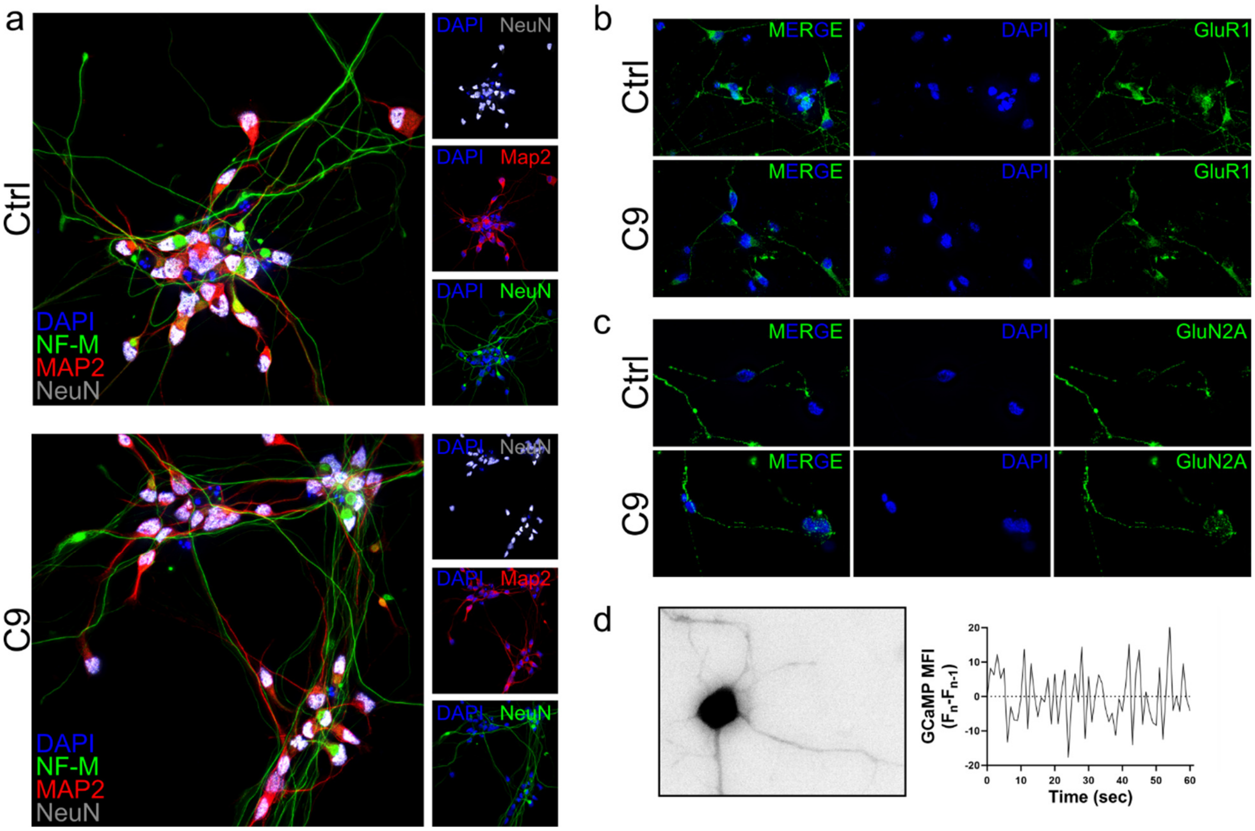


**Supplementary Figure 3. iPSC-derived cortical neuron differentiation. a.** Representative images of iPSC-derived cortical neurons (i$^3$CNs) 2 weeks after differentiation stained with pan-neuronal neurofilament (NF-M, *green*), MAP2 (*red*), and NeuN (*grays*). Similar expression levels of all markers were observed between control and C9 neurons. **b-c.** Two-week-old C9 and control i$^3$CNs display similar surface expression of glutamate receptors GluR1 (*b, green*) and GluN2A (*c, green*). **d.** At 14 days, i3CNs are able to spontaneously fire action potentials, as detected by changes in the genetically encoded calcium sensor GCaMP6f. A representative image of a GCaMP-positive neuron and the changes in fluorescence levels of GCaMP over a 60 second video is shown.

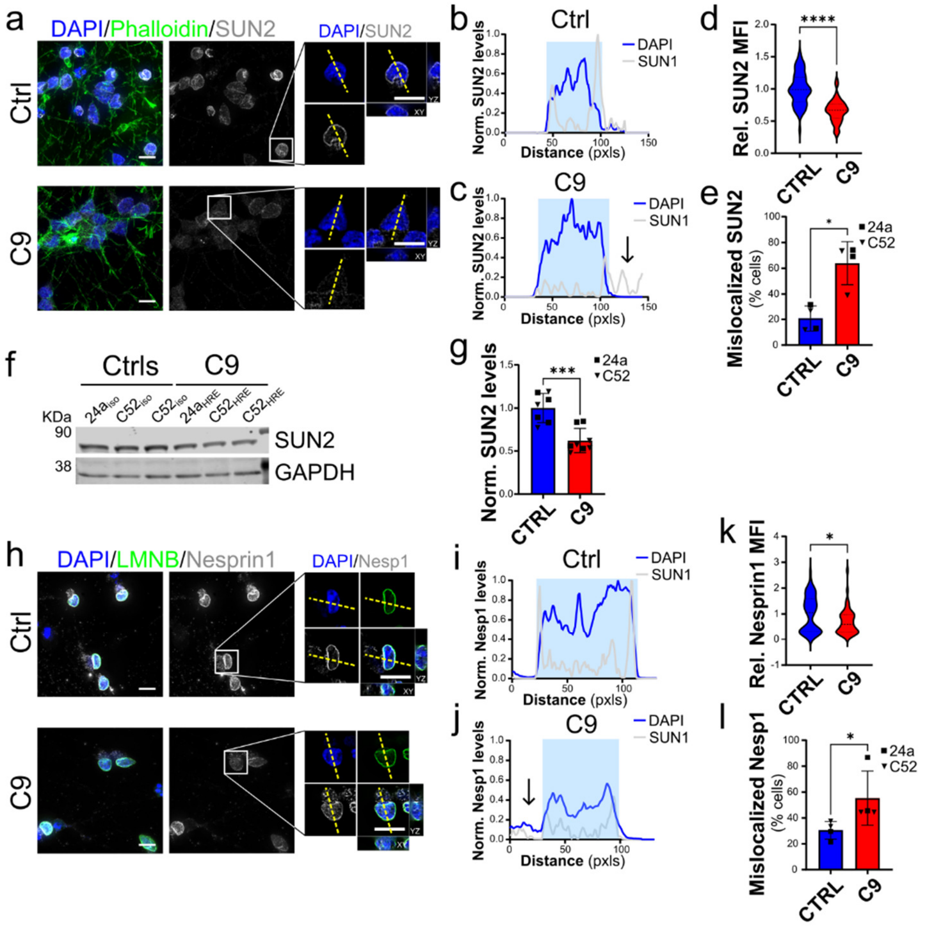


**Supplementary Figure 4. SUN2 and Nesprin1 alterations in C9-ALS i$^3$CNs. a.** Representative images of SUN2 (*grays*) in C9-ALS and control i$^3$CNs. DAPI (*blue*) identified the nuclei, while Phalloidin (*green*) labeled the actin cytoskeleton. The white boxes indicate the neurons enlarged in the panels on the right. **b-d.** Plots of line profiles of SUN2 and DAPI intensities show a higher frequency of SUN2 mislocalization to either the nucleoplasm or cytoplasm (arrows), as quantified in *d*. The yellow dashed lines in *a* indicate the lines used for the profile plots (Mann-Whitney *t* test, n= 4, **p*<0.05). **e**. The quantification of the nuclear mean fluorescence intensity (MFI) of SUN2 in C9 i$^3$CNs shows a significant reduction in its abundance relative to isogenic controls (Student's *t* test, n=60 CTRL and C9 neurons respectively from 4 independent differentiations, *****p*<0.0001). **f-g.** Representative western blot (WB) and quantification of SUN2 levels relative to GAPDH expression shows a significative reduction of total SUN2 protein levels in C9 lines compared to isogenic counterparts (n=7 and 8 independent experiments for both 24a and C52 C9 and iso neurons; Student's *t* test, *** *p* < 0.001). **h.** Representative images of Nesprin1 (Nesp1) staining in i$^3$CNs from C9 and Ctrl iPSC lines. DAPI (*blue*) identified the nuclei, while LaminB (LMNB, *green*) labeled the nuclear lamina. The white boxes indicate the neurons enlarged in the panels on the right. **i-k.** Plots of line profiles of Nesprin1 and DAPI intensities normalized to the max intensity in control cells shows an increase in the frequency of Nesprin1 mislocalization to either the nucleoplasm or cytoplasm (arrows), quantified in *k*. The yellow dashed lines in *h* indicate the lines used for the profile plots (Mann-Whitney *t* test, n= 4, **p*<0.05). **l.** The quantification of the Nesprin1 relative nuclear MFI shows a significant reduction in its abundance in C9 i$^3$CNs compared to isogenic controls (Mann-Whitney *t* test, n=58 and 48 for CTRL and C9 neurons from 4 independent differentiations, **p* < 0.05). Scale bars: 20µm in main panels, 10µm in zoomed-in images. For all, bars are mean and SEM, while violin plots show the distribution of the data with dashed lines indicating median and quartiles.

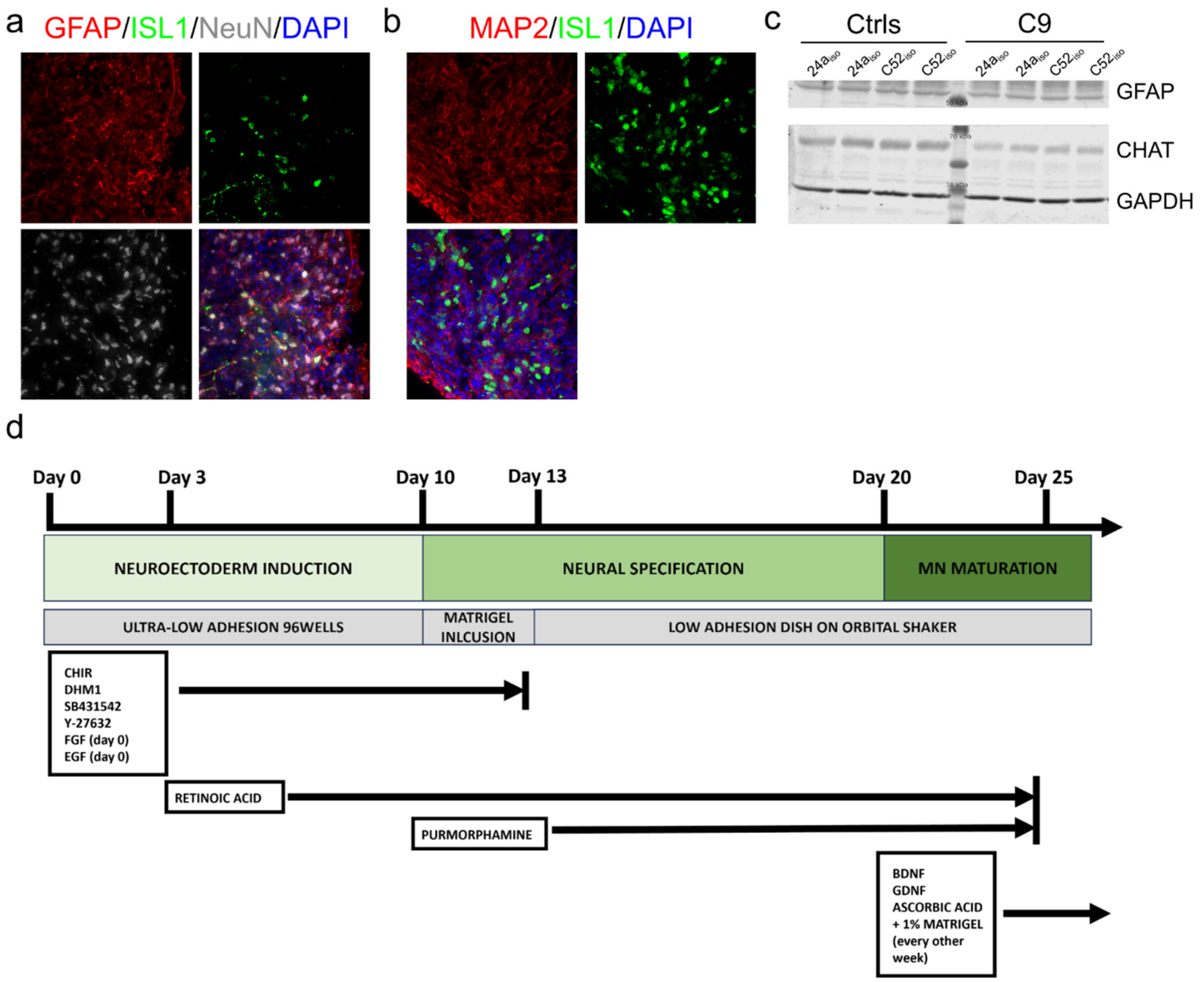


**Supplementary Figure 5. iPSC-derived spinal cord organoids differentiation. a-b.** Representative images of 50-day-old spinal organoids. Motor neurons were identified via Islet1 staining (*green* in *a* and *b*), while astrocytes were labeled with GFAP antibody (*red* in *a*). NeuN (*grays* in *a*) and MAP2 (*red* in *b*) were used as pan-neuronal markers. DAPI (*blue*) labeled the cells' nuclei. **c.** Western blot analysis of CHAT and GFAP show similar levels of expression in 50-day-old organoids from all isogenic and mutant lines used (i.e. 24a and C52). **d.** Schematic representation of organoid differentiation protocol.

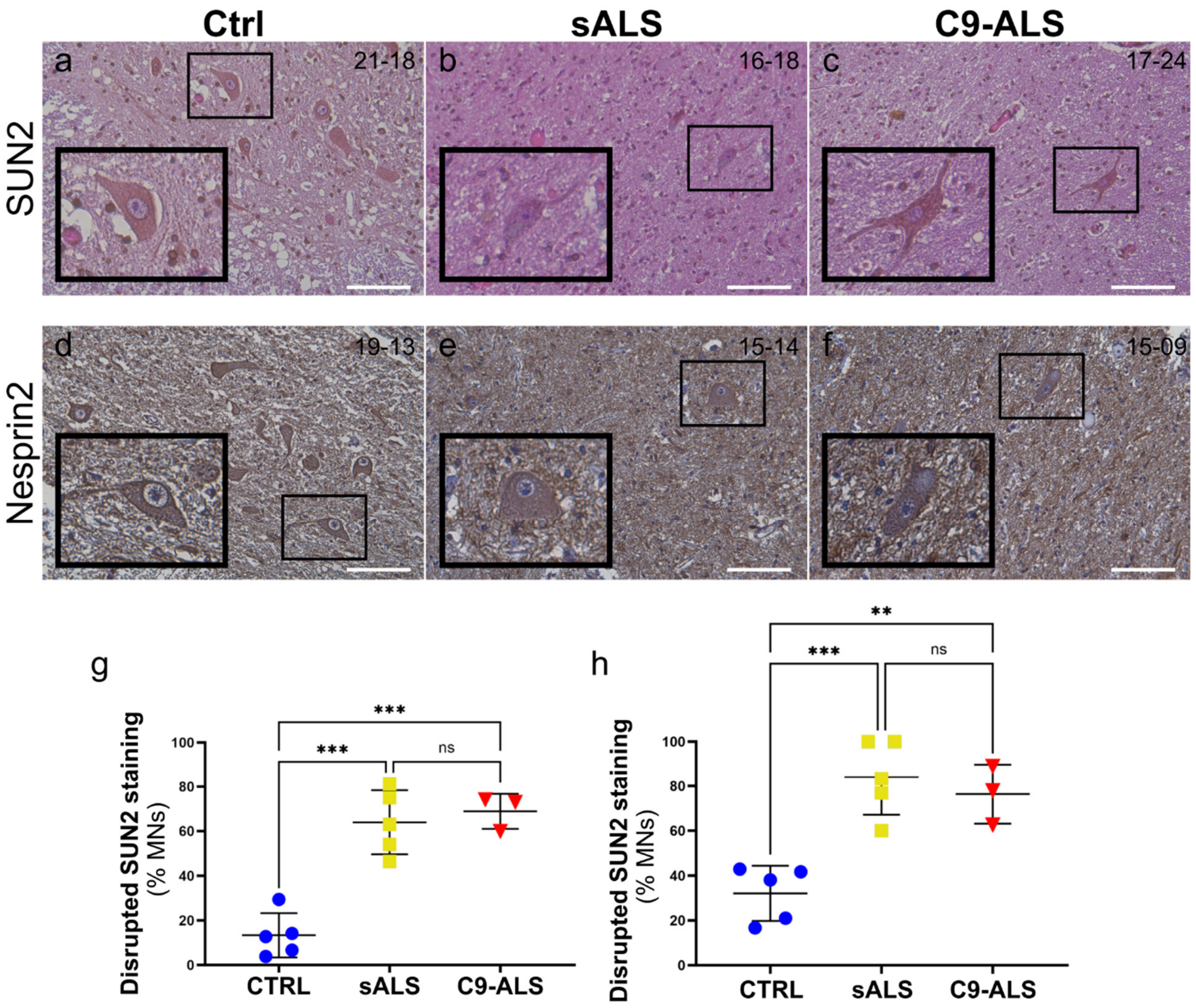


**Supplementary Figure 6. SUN2 and Nesprin2 disruption in sALS and C9-ALS spinal cord *postmortem* biopsies. a-f.** Spinal cord sections from control (*a, d*), sALS (*b, e*) and C9-ALS (*c, f*) patients were stained with antibodies specific for SUN2 (*a-c*) and Nesprin2 (*d-f*). Hematoxylin and eosin counterstains were used to identify the nucleus and cytoplasm, respectively. The black boxes identify the motor neurons enlarged in the insets. Scale bars: 100μm. **g-h.** The frequency of disrupted NE staining for both SUN2 (*g*) and Nesprin2 (*h*) was quantified blindly in at least two sections from each patient's tissue. A significant increase in the percentage of cells with disrupted staining was observed in both sALS and C9-ALS spinal cords compared to controls. Each dot represents the mean of at least two sections for each case, horizontal lines show mean and standard deviation (one-way ANOVA with Tukey *post hoc* test, n=5, 5, and 3, ***$p$<0.001, ns=not significant).

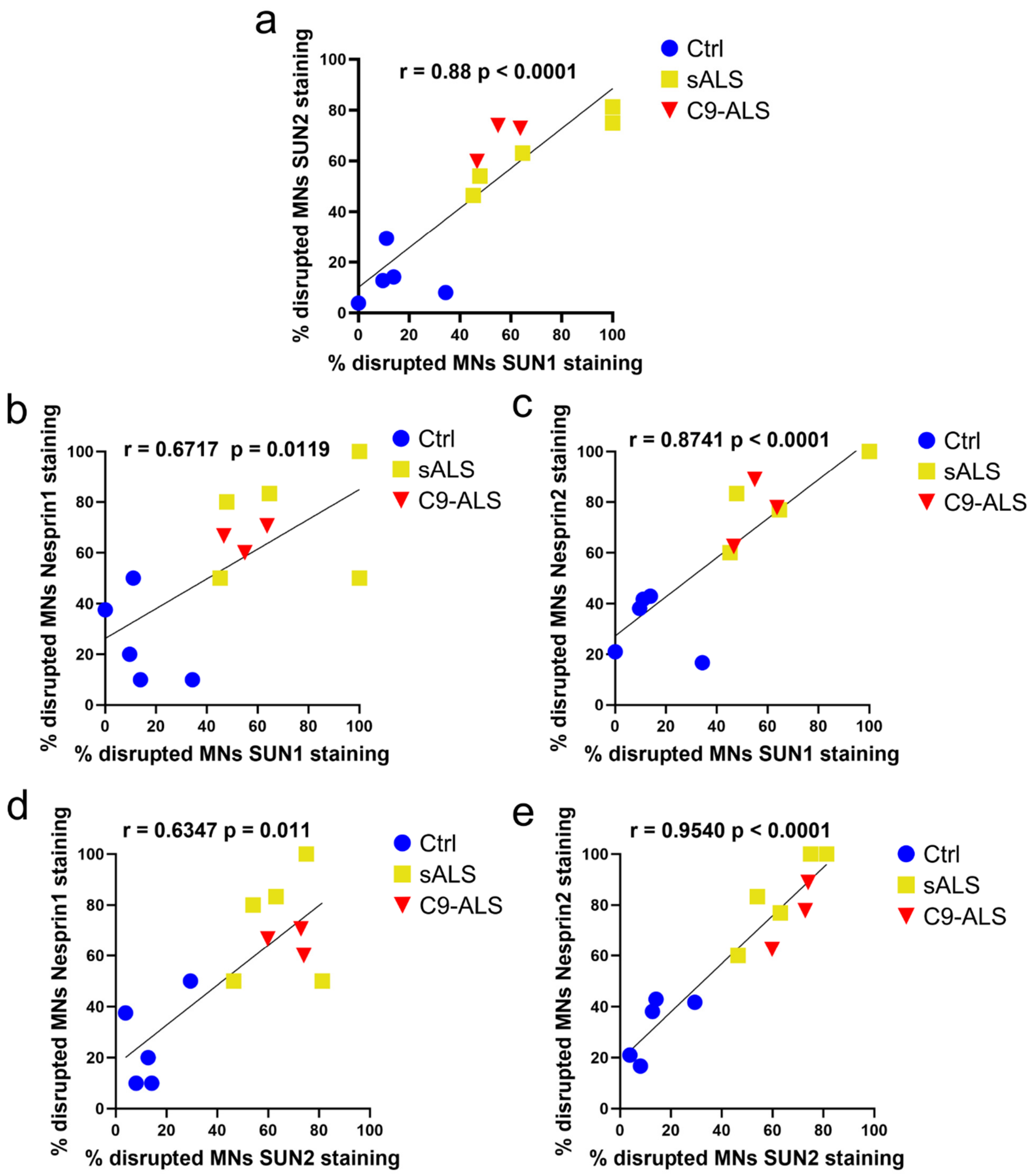


**Supplementary Figure 7. Correlative analysis of LINC protein disruption in spinal cord biopsies.** Linear regression analyses show a high degree of correlation between the levels of alterations in the staining profiles of SUN1 and SUN2 (***a***), Nesprin1 (***b***) and Nesprin2 (***c***), as well as SUN2 and Nesprin1 (***d***) and Nesprin2 (***e***).

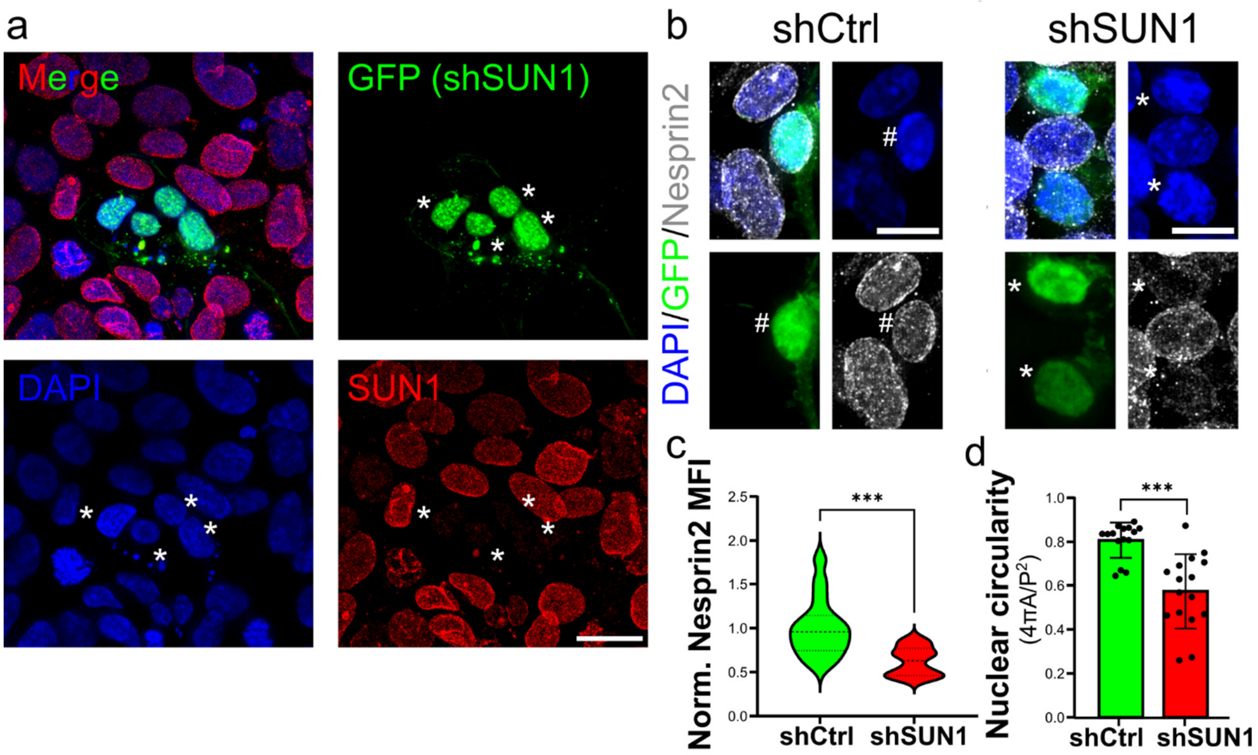


**Supplementary Figure 8. SUN1 is fundamental for Nesprin2 nuclear envelope localization. a.** Representative images of HEK293 cells transfected with a shRNA (shSUN1) to knock down SUN1 (shSUN1). GFP (*green*) is co-expressed with the shRNA and was used to identify the transfected cells. GFP-positive cells display an almost complete absence of SUN1 immunoreactivity (*red*) compared to un-transfected cells. **b.** Representative images and quantification of Nesprin2 levels (*grays*) in GFP-positive cells (*green*) that express shSUN1 (right panel, * identify transfected cells) or a non-silencing control shRNA (shCtrl, left panel, # identify a transfected cell). **c.** A significant reduction in Nesprin2 nuclear levels was observed in shSUN1 transfected cells compared to controls (Student's *t* test, n= 16 from 3 independent experiments, ***$p$<0.001). d. The quantification of nuclear circularity shows that reduction in the nuclear levels of LINC proteins causes morphological alterations to the nucleus (Mann-Whitney *t* test, n=15 and 16, ***$p$<0.001). DAPI (*blue*) was used to label the cells' nuclei. Scale bars: 20μm in main panels, 10μm in zoomed-in images.

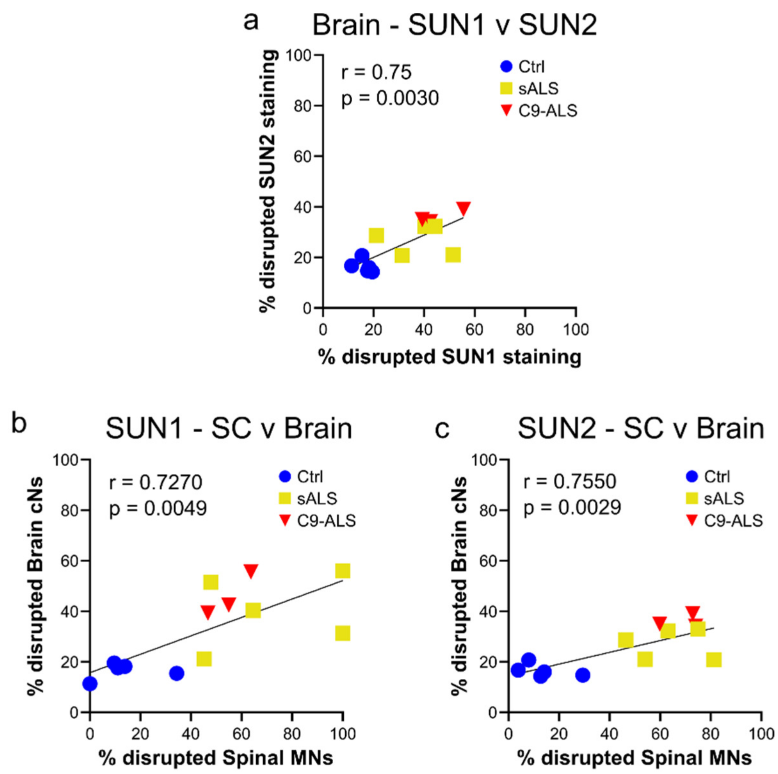


**Supplementary Figure 9. Correlative analysis of SUN proteins disruption. a.** Linear regression analyses show a high degree of correlation between the levels of alterations in the staining profiles of SUN1 and SUN2 in brain neurons. **b-c.** Scatterplots show the correlation between the degree of SUN1 (*b*) or SUN2 (*c*) disruption along the neural axis (i.e. spinal cord (SC) versus brain). Linear regression analyses were performed to support significance of the correlation.

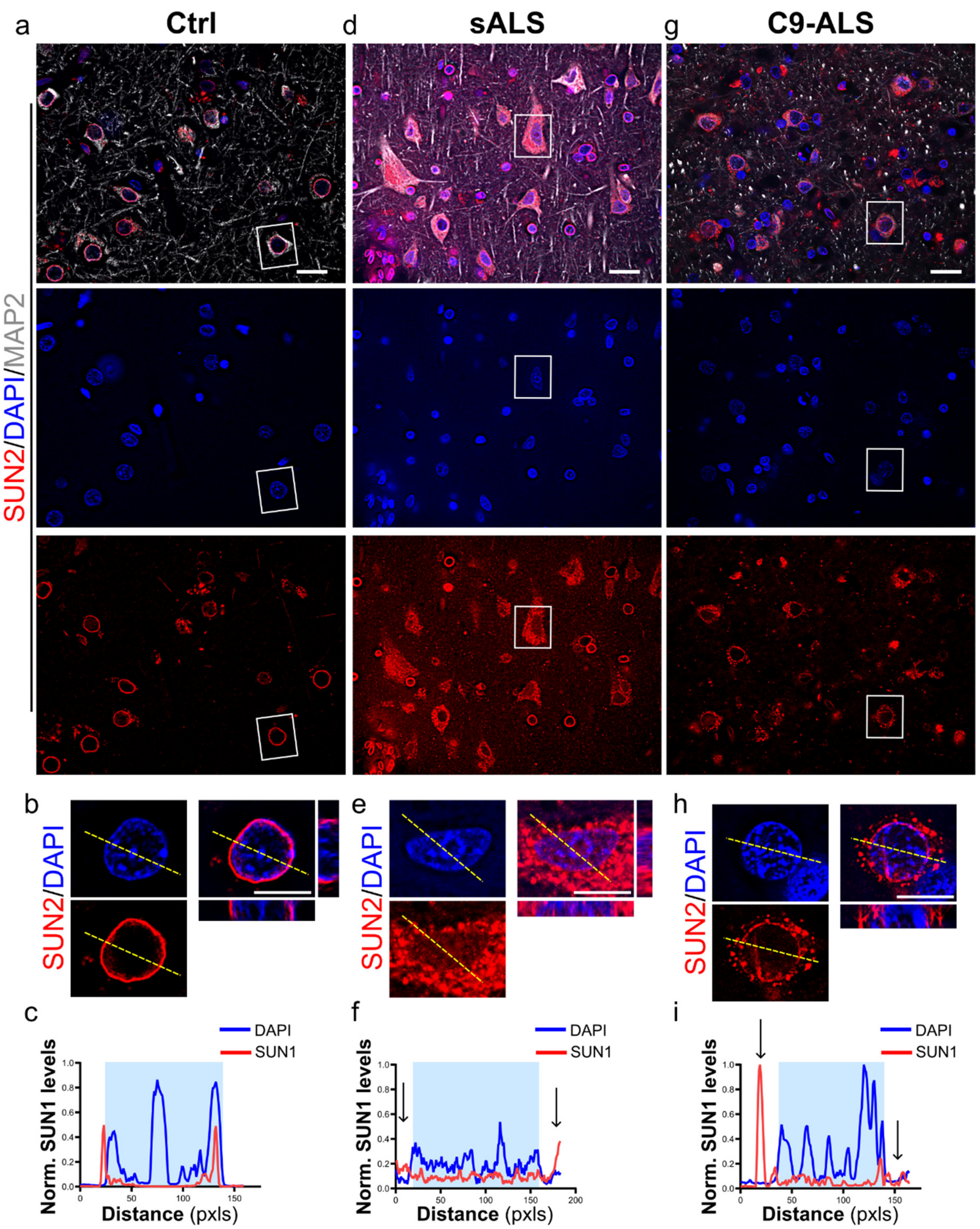


**Supplementary Figure 10. SUN2 mislocalization in the cytoplasm in cortical neurons of sporadic and C9-ALS patients.** Representative images of motor cortex sections from control (***a, b***), sALS (***d, e***) and C9-ALS (***g, h***) patients stained with antibodies specific for SUN2 (*red*) and MAP2 (*grays*). DAPI (*blue*) was used to label nuclei. White boxes in *a*, *d*, and *g* identify neurons enlarged in *b*, *e*, and *h*. Line profile plots show the marked difference of SUN2 distribution in sALS (***f***) and C9-ALS (***i***) cortical neurons compared to controls (***c***). The yellow dashed lines in *a* indicate the lines used for the profile plots. Scale bars: Scale bars: 20μm in main panels, 10μm in zoomed-in images.

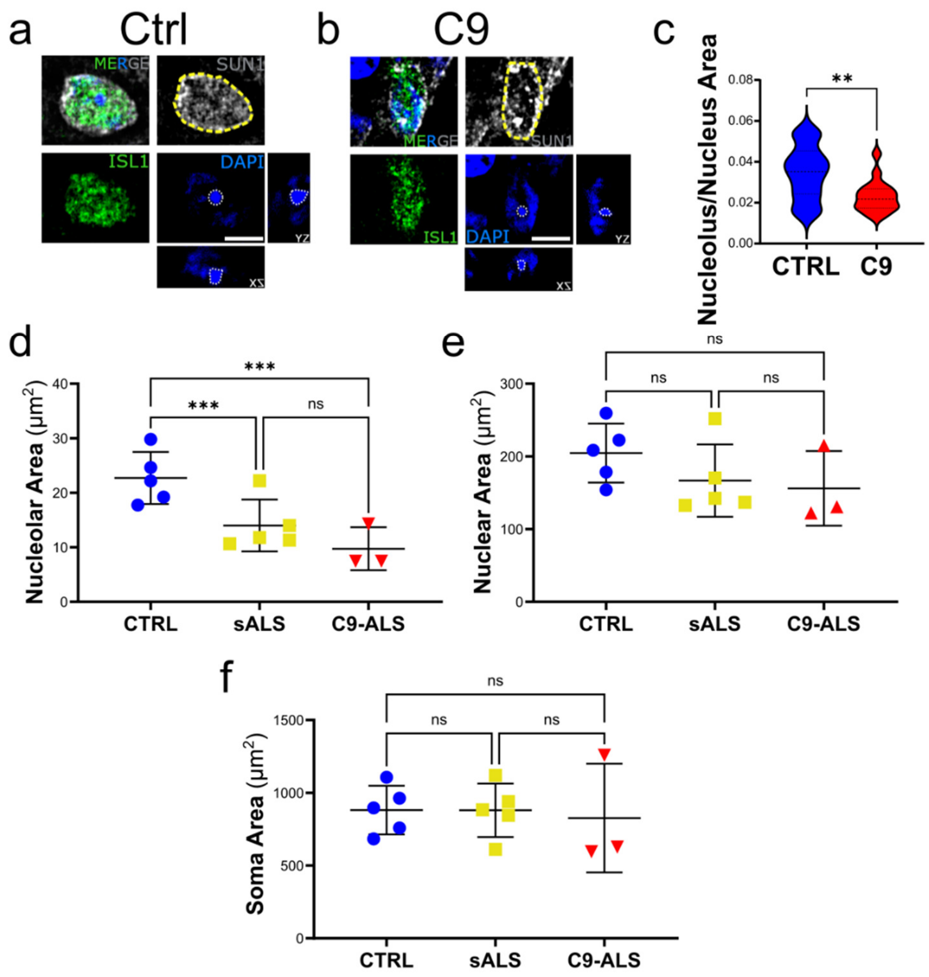


**Supplementary Figure 11. Loss of SUN1 correlates with altered nuclear and nucleolar size. a-b.** Representative images of SUN1 (*grays*) staining in Islet1-positive iMNs (*green*) in spinal organoid from control (*a*) and C9 mutant (*b*) iPSCs. Yellow dashed lines in SUN1 panels outline the nuclear envelope, while white dashed lines in DAPI panels identify the nucleoli. Scale bars: 10μm. **c.** Quantification of nucleolar area normalized to nuclear size shows a significant reduction in C9 iMNs compared to control cells (Student's *t* test, n=23 and 17, ***p*<0.001). **d-f.** Measurement of the nucleolar, nuclear and somatic size shows a significant reduction in the absolute area of the nucleoli of sALS and C9-ALS spinal cord motor neurons compared to controls, and a trend toward smaller nuclei. No major difference was observed in the average size of MN somas (one-way ANOVA, n=5, 5, 3. ****p*<0.001, ns= not significant).

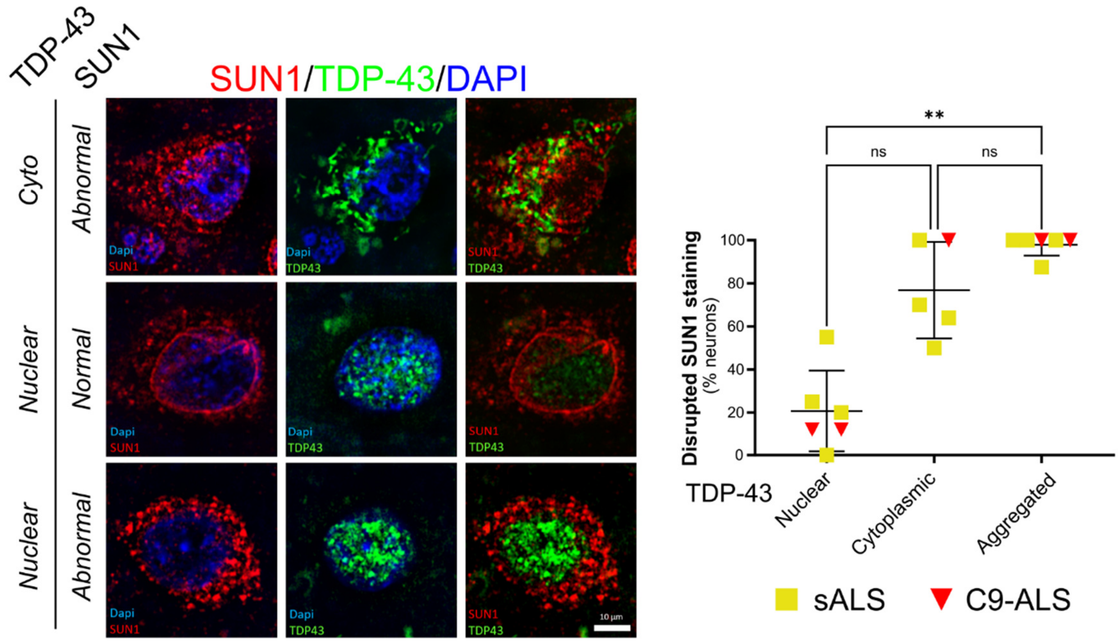


**Supplementary Figure 12. TDP43 aggregation strongly associates with SUN1 disruption.** Representative images of cortical neurons from ALS cases presenting aggregated TDP43 (*green*) and disrupted SUN1 (*red, top*), nuclear TDP43 and normal SUN1 (*middle*), and nuclear TDP43 but disrupted SUN1 staining (*bottom*). DAPI (blue) labeled the nuclei. The frequency of SUN1 staining alterations was quantified in neurons according to their TDP-43 state in 4 sALS and 2 C9-ALS cases. In every of the analyzed cases, cNs presenting TDP43 aggregation also displayed SUN1 disruption. However, SUN1 disruption was also occurring in ALS cNs even in absence of TDP43 pathology (one-way ANOVA with Tukey post hoc test, n=5, ***p*<0.01, ns=not significant). Scale bar: 10μm.

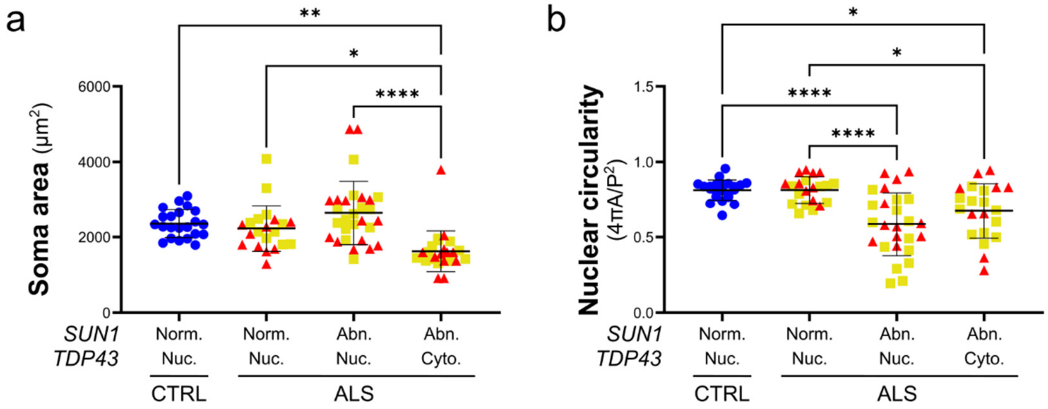


**Supplementary Figure 13. Morphological changes in cortical neurons depend on both TDP-43 and SUN1.** Quantitative analysis of cell soma area (***a***) and nuclear circularity (***b***) in cortical neurons categorized based on the presence of nuclear (Nuc.) or cytoplasmic (Cyto.) TDP-43 and normal (Norm.) or abnormal (Abn.) SUN1 staining shows that cytoplasmic accumulation of TDP-43 alone is sufficient to cause a significant reduction in the size of the cell soma, while loss of SUN1 from the NE is a main driver of changes to nuclear shape (one-way ANOVA with Tukey post *hoc* test, n=22, 21 ,29 ,24 in *a*, n=22, 21, 26, and 21 in *b*,  **p*<0.05, ***p*<0.01, *****p*<0.0001).

| Name | Line ID | Genotype | Reference/Source |
|---|---|---|---|
| $24a_{HRE}$ | 180906.4a | C9 6 kb HRE | Ababneh et al., 2020 |
| $24a_{iso}$ | 180906.2a | Isogenic | Ababneh et al., 2020 |
| $C52_{HRE}$ | CS52iALS-C9n6 | C9 6-8 kb HRE | Cedars-Sinai |
| $C52_{iso}$ | CS52iALS-C9n6.ISOC3 | Isogenic | Cedars-Sinai |

**Supplementary Table 1.** List of iPSC lines used in the study with the name used in this study, the official line ID, and the relevant source.

| Case # | Sex | Age at Death | Cause of death | PMI (hrs.) |
|---|---|---|---|---|
| GBB-20-16 | Male | 71 yrs. | Metastatic liver cancer, gastric adenocarcinoma, acute tubular necrosis | 24 |
| GBB-19-13 | Male | 75 yrs. | Renal cell carcinoma, pulmonary thromboembolism | 8 |
| GO-21-18578 GBB-21-27 | Female | 82 yrs. | Diffuse large cell lymphoma, hypertension | 27 |
| GBB-17-22 | Male | 70 yrs. | Metastatic cecal carcinoma | 91 |
| GBB-21-03 | Male | 72 yrs. | Hypertension, chronic kidney disease, bronchiectasis, acute hypoxic respiratory failure | 47 |

**Supplementary Table 2. Clinical and pathological information of non-neurological control cases.** PMI, postmortem interval.

| Case # | Sex | Age at Death | Age at Onset | Site of Onset | Disease duration | PMI (hrs.) | Clinical Diagnosis | Mutation |
|---|---|---|---|---|---|---|---|---|
| GWF15-09 | Female | 61 yrs. | 59 yrs. | n.a. | 17 mos. | 5.5 | ALS | C9 HRE |
| GWF15-10 | Female | 63 yrs. | 61 yrs. | Limb | 24 mos. | 37.5 | ALS | None |
| GWF15-14 | Male | 74 yrs. | n.a. | n.a. | n.a. | 3.5 | ALS | None |
| GWF16-18 | Female | 60 yrs. | 57 yrs. | Limb | 32 mos. | 4 | ALS | None |
| GWF16-20 | Female | 63 yrs. | 60 yrs. | Bulbar/limb | 37 mos. | 4 | ALS | None |
| GWF17-24 | Female | 51 yrs. | 47 yrs. | Bulbar | 60 mos. | 7 | ALS | C9 HRE |
| GWF18-31 | Female | 62 yrs. | 60 yrs. | Limb | 20 mos. | 3.5 | ALS-FTD | C9 HRE |
| GWF19-35 | Male | 40 yrs. | 32 yrs. | Limb | 98 mos. | 6 | ALS | None |

**Supplementary Table 3. Clinical and Pathological information of ALS cases.** PMI, postmortem interval; n.a., not available.

| Antibodies | Company | Identifier | Catalog number | Working Dilution |
|---|---|---|---|---|
| SUN1 | Novus Biologicals | | NBP1-87396 | 1:200 (IF/IHC) 1:1000 (WB) |
| SUN2 | Proteintech | AB_2880906 | 27556-1-AP | 1:200 (IF/IHC) 1:1000 (WB) |
| NESPRIN1 | Novus Biologicals | | NBP1-89349 | 1:200 (IF/IHC) |
| NESPRIN2 | Novus Biologicals | | NBP1-84190 | 1:200 (IF/IHC) |
| MAP2 | Invitrogen | AB_2138189 | PA1-16751 | 1:1000 (IF) |
| ISL1 | DHSB | AB_528315 | 40.2D6 | 0.2-0.5 ug/ml (IF) |
| Alexa Fluor™ 488 Phalloidin | Thermo scientific | | A12379 | 1:50 |
| LAMIN-B | Proteintech | AB_11232208 | 66095-1-Ig | 1:500 (IF) |
| TDP43 | Proteintech | AB_615042 | 10782-2-AP | 1:1000 (IF) |
| GFAP | HelloBio | | HB6406 | 1:2000 (IF) 1:5000 (WB) |
| GluR1 | NeuroMab | AB_2315840 | SKU: 75-327 | 1:500 (IF) |
| NR2A | NeuroMab | AB_2315842 | SKU: 75-288 | 1:500 (IF) |
| Histone-H3 | Proteintech | AB_2716755 | 17168-1-AP | 1:2000 (WB) |
| NeuN | Invitrogen | AB_2633050 | 702022 | 1:500 (IF) |
| CHAT | Millipore | | MAB305 | 1:500 (WB) |
| PAX6 | DHSB | AB_528427 | PAX6 | 2-5 ug/ml |
| OLIG2 | Millipore | | AB9610 | 1:200 (IF) |
| GAPDH | Proteintech | AB_2107436 | 600004-1-1g | 1:5000 |

| Chemicals | Company | Catalog number | Working Concentration |
|---|---|---|---|
| Y-27632 | SellChem | S1049 | 10 µM |
| FGF | Corning | 354060 | 10 ng/mL |
| EGF | Corning | 354052 | 20 ng/mL |
| CHIR99021 | Tocris | 4423 | 3 µM |
| DMH1 | Tocris | 4126/10 | 2 µM |
| SB431542 | Tocris | 1614/1 | 2 µM |
| Retinoic acid | Thermo scientific | 207341000 | 0.5 µM |
| Purmorphamine | Tocris | 4551 | 1 µM |
| Ascorbic Acid | Sigma | A4544 | 0.1 mM |
| BDNF | Proteintech | HZ-1335 | 20 ng/mL |
| GDNF | Proteintech | HZ-1311 | 10 ng/mL |
| Matrigel Matrix for Organoid Culture | Corning | 356255 | |

**Supplementary Table 4.** Detailed information on reagents and resources used.
